# eTRex Reveals Oncogenic Transcriptional Regulatory Programs Across Human Cancers

**DOI:** 10.64898/2026.01.16.699950

**Authors:** Zeyu Lu, Yuqiu Yang, Qiang Zheng, Feng Gao, Lin Xu, Xinlei Wang

## Abstract

Transcriptional regulators (TRs) are essential proteins that regulate gene transcription, and their dysregulation defines the oncogenic transcriptional regulatory programs that drive tumor-specific gene expression. Existing pan-cancer resources summarize these programs by aggregating signals across individual datasets, neglecting the context-specific features that capture the diversity of transcriptional regulation underlying cancer heterogeneity. By developing a variational Bayesian hierarchical model named eTRex (**e**pigenomics-based **T**ranscriptional **R**egulator **ex**plorer) and applying it to 4,819 cancer-related ATAC-seq datasets, we provide a comprehensive pan-cancer atlas of functional TR profiles that preserve the context-specific features of each dataset. This study reveals both common regulators across diverse malignancies and those with highly specific roles. We extensively validated these findings using independent CRISPR/Cas9 screening, mutation, and transcriptomic datasets. Collectively, this pan-cancer atlas of functional TR profiles represents a comprehensive, biologically interpretable resource for uncovering transcriptional regulatory programs, identifying biomarkers, prioritizing therapeutic targets in oncology, and is freely accessible through an interactive web portal.

## Introduction

Transcriptional regulation is a fundamental biological process that controls gene expression^1^. This intricate process depends on the coordinated actions of diverse transcriptional regulators (TRs)—including transcription factors, co-factors, chromatin remodelers, and other regulatory molecules^2–4^. Dysregulation of TRs can drive tumor initiation, progression, and therapeutic resistance^5–7^. Although more than 2,000 TRs have been catalogued, their functional roles vary markedly across tumor types and cellular contexts. For example, FOXA1 functions as a cofactor of the androgen receptor that shapes enhancer selection and whose dysregulation drives oncogenic reprogramming of AR signaling in prostate cancer^8^, whereas in luminal breast cancer it acts as a pioneer factor that stabilizes estrogen-dependent transcription and differentiated luminal identity^9^. Therefore, understanding the context-specific functions of TRs remains a major focus in biomedical research.

Experimental assays such as ChIP-seq, CUT&Tag, and CRISPR perturbations are the gold standard for assessing TR function^10–12^, yet they are costly, labor-intensive, and require high-quality antibodies or genetic tools. As a result, systematically characterizing the functional roles of all 2,000 TRs across diverse biological contexts is infeasible. Indeed, even well-resourced laboratories can only investigate a limited subset of regulators relevant to their specific research interests. To address this challenge, researchers have increasingly turned to the publicly available omics data, which contain latent regulatory information that can be computationally analyzed to infer TR activity and function^13–15^. Leveraging these resources enables the identification and prioritization of TRs that are most likely to exert key regulatory roles. Statistical and machine learning–based computational frameworks have therefore emerged as powerful and scalable tools to prioritize functional TRs, offering a data-driven complement to experimental validation and mechanistic studies^16–20^.

However, most applications of these computational frameworks have been tailored to one research topic, such as individual cancer types, experimental conditions, or disease cohorts. For instance, computational methods have been used to identify upstream regulators of NUTF2 target genes in endocrine-resistant breast cancer^21^, or to predict functional TRs associated with ARID1A loss in prostate tumorigenesis^22^. While such studies yield valuable insights, they provide only a limited view of transcriptional regulation across malignancies. In contrast, a pan-cancer perspective can bridge these isolated efforts by systematically analyzing the function of TRs across diverse tumor lineages and conditions^23–26^. Such integrative analysis enables the identification of both shared and specific regulators, reveals conserved oncogenic circuits, and supports the discovery of regulatory biomarkers and therapeutic targets.

Despite this conceptual advance, practical implementations remain limited. To establish a pan-cancer atlas of functional TR profiles, we systematically curated 4,819 cancer-related ATAC-seq datasets, each profiling a unique chromatin accessibility landscape. Given that accessible chromatin regions are prerequisites for TR binding, these datasets serve as query data for functional TR inference. Notably, several recent studies also attempted to define a pan-cancer atlas of functional TRs^23–25^, but often at a much coarser scale that overlooks critical regulatory differences between cancer subtypes or individual cell lines. For example, one study aggregated all ATAC-seq datasets belonging to the same TCGA cancer type to create a single chromatin accessibility landscape per type^24^. When we compared these aggregated landscapes with the profiles of the 4,819 individual ATAC-seq datasets collected in this study, we observed substantial variability across datasets, suggesting that context-specific epigenomic features may be obscured (**Supplementary Figure 1**). By contrast, this study aims to produce an atlas that fully preserves such context-specific information.

In addition, although numerous computational frameworks have been proposed, accurate identification of functional TRs still presents significant challenges, as discussed by our recent comprehensive review^27^. Methods such as ChIP-Atlas^13^, WhichTF^17^, BART^18^, i-cisTarget^19^, and HOMER^20^ rely either on linear genomic distance to infer regulatory potential or on TR binding motifs as references, which may fail to capture context-specific TR binding sites. Moreover, these methods often lack a formal treatment of both within- and between-TR heterogeneity arising from multiple ChIP-seq profiles associated with the same TR as well as from different TRs, increasing the risk of overfitting. Our earlier method BIT provided an initial step towards addressing these challenges^28^. However, its dependence on Gibbs sampling via Markov chain Monte Carlo (MCMC) limits its computational efficiency for large-scale analysis.

With the rapid growth of epigenomics technologies and the increasing availability of thousands of TR-related profiles across diverse cancer types, scalable computational methods have become indispensable for practical and timely TR discovery. Thus, to allow accurate and scalable identification of pan-cancer functional TRs, we introduce eTRex (**e**pigenomics-based **T**ranscriptional **R**egulator **ex**plorer), a novel framework based on mean-field variational inference^29^ with a coordinate ascent algorithm^29^ that admits closed-form updates at each iteration. Given a query ATAC-seq dataset, eTRex evaluates genome-wide consistency between its accessible chromatin regions and TR binding sites from an internal ChIP-seq reference database to assign a eTRex score to each candidate regulator, which indicates its regulatory influence within the corresponding biological context. eTRex delivers substantial advances in scalability and computational efficiency while achieving the top-level accuracy in TR identification, representing a significant improvement over existing approaches.

We applied eTRex to all 4,819 ATAC-seq datasets to generate dataset-specific functional TR profiles, which together constitute a pan-cancer atlas of TR activity. Unlike previous integrative studies, eTRex retains the fine-grained variability inherent to individual ATAC-seq datasets, thereby capturing the nuanced regulatory programs that define specific cell lines, tissues, or disease states. Furthermore, eTRex employs mean reciprocal rank fusion (MRRF) to flexibly aggregate individual TR profiles into any user-defined research tasks. Through this design, eTRex preserves context-specific regulatory information at the individual dataset level and enables flexible aggregation for high-level analyses. This resource identifies ubiquitous regulators that operate across multiple malignancies as well as highly specific TRs that define distinct cancer subtypes. In addition, it provides a scalable and reusable framework that enables researchers to interrogate TR activity in virtually any biological context.

Finally, we cross-validated the eTRex-identified regulators using complementary evidence from cellular viability, gene activity, and somatic mutation frequency, providing additional supports for their critical functional roles. Overall, this work contributes: (1) a pan-cancer atlas of functional TR profiles, preserving the context-specific information and offering flexible re-aggregation across user-defined research tasks; (2) a novel variational Bayesian hierarchical model enabling scalable identification of functional TRs across cancers; (3) comprehensive validation of eTRex-identified regulators; (4) systematic discovery of cancer-type- and subtype-specific TRs; and (5) an interactive web portal to facilitate community access and analysis (https://e-trex.org).

## Results

### Development and validation of the eTRex method

Transcriptional regulators generally exert their function by binding to regions of open chromatin, which are accessible for protein–DNA interactions^30,31^. These accessible regions can be profiled using epigenomic assays such as ATAC-seq^32^, DNase-seq^33^, or FAIRE-seq^34^. Among these, ATAC-seq has become the technique of choice because it requires minimal input material, offers high resolution and genome-wide coverage, and is rapid, cost-effective, and easy to implement across diverse biological systems^35^. In parallel, the binding sites of an individual TRs can be directly mapped using ChIP-seq, the gold-standard approach for this purpose^36^. eTRex thus sourced 10,140 human TR ChIP-seq datasets from GTRD^14^ to construct a comprehensive database covering 988 human TRs.

First, to systematically link chromatin accessibility to TR binding sites, eTRex encodes genome-wide chromatin-accessible regions from a query ATAC-seq dataset and TR binding profiles from reference ChIP-seq datasets into binary vectors of the same length (**Figure 1a**). To achieve this, the human genome is partitioned into consecutive, non-overlapping bins of fixed length (e.g., 1000 base pairs). Each bin is then annotated with a binary label: “1” if it overlaps with at least one peak (i.e., a region identified as significantly enriched for chromatin accessibility in ATAC-seq or TR binding in ChIP-seq), and “0” otherwise. This step transforms irregular peaks into compact, standardized binary vectors. By applying this procedure uniformly across all datasets, the entire TR ChIP-seq reference database is preprocessed into a library of fixed-length binary vectors, drastically reducing storage requirements and enabling rapid comparison.

**Figure 1.**
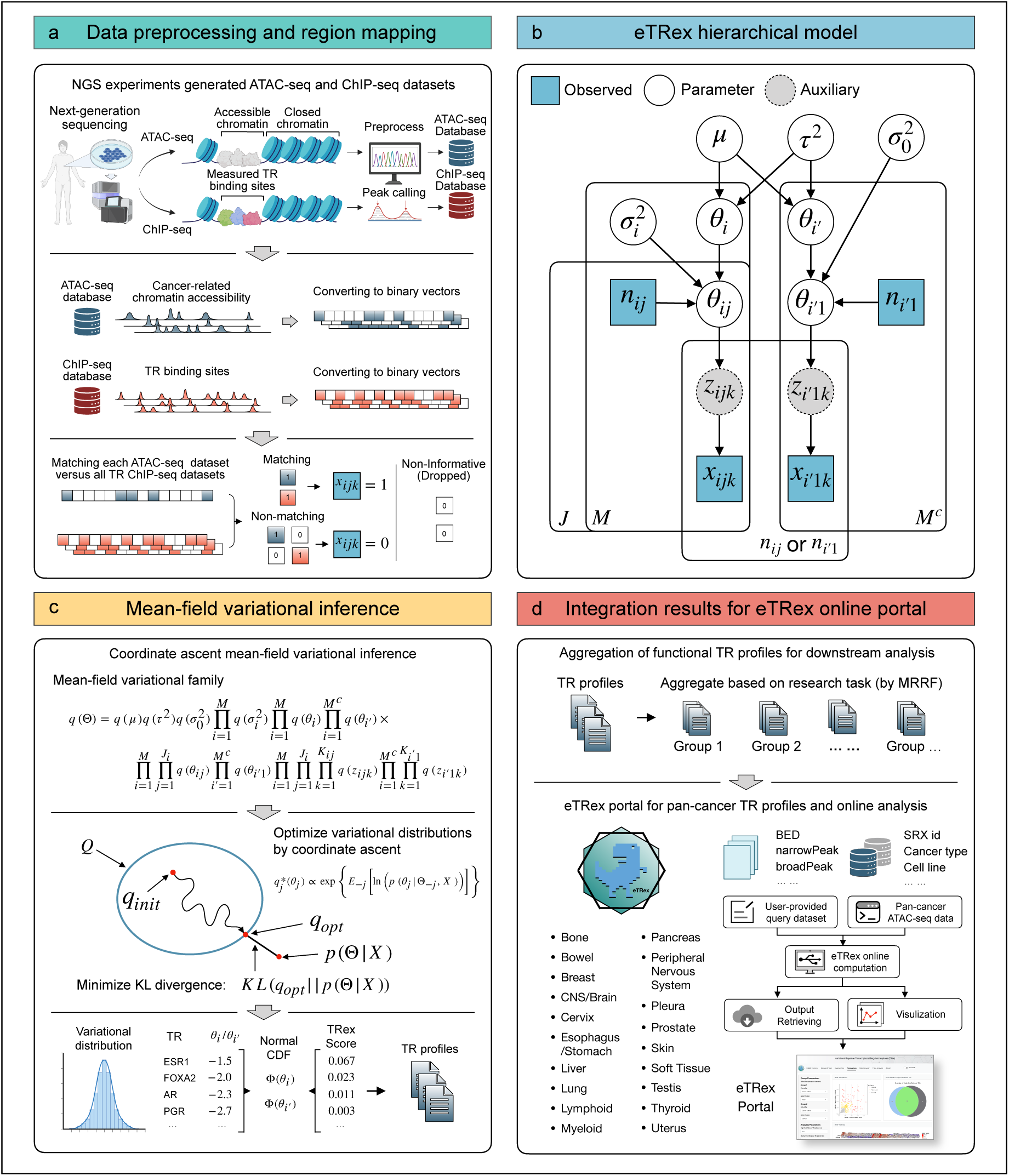
overview of the eTRex model. (a). Next-generation sequencing experiments can measure chromatin accessibility profiles (e.g., ATAC-seq) and transcriptional regulator binding sites (ChIP-seq) across different biological conditions. Pan-cancer ATAC-seq datasets are used as input. TR ChIP-seq datasets are used as reference. These datasets are uniformly converted to binary vectors. The converted ATAC-seq vectors are compared with reference ChIP-seq vectors. (b). eTRex hierarchical model leverages a probit regression model and a latent data augmentation strategy to integrate information across multiple TRs and TR ChIP-seq datasets. (c). eTRex uses mean-field variational inference with coordinate ascent algorithm to estimate *θ*_*i*_ and *θ*_*i*′_ that represent TR-level consistency. Scores are computed using the standard normal cumulative distribution function to transform estimated *θ*_*i*_ and *θ*_*i*′_ into a probability scale. (d). Pan-cancer TR profiles are saved on eTRex online portal for easy access. The online portal also implemented eTRex for online analysis.

When a query ATAC-seq dataset is submitted for analysis, eTRex processes it using the exact same binning step, generating a query binary vector that mirrors the structure of the reference library. Identification of functional TRs is then performed by counting the overlap between the query vector and each reference vector. Crucially, because both query and reference data are already represented in compatible binary form, eTRex avoids on-the-fly genomic interval arithmetic or dynamic peak overlap calculations. This precomputation strategy not only accelerates inference (scaling linearly with the number of reference TRs) but also ensures reproducibility and consistency across analyses.

With this framework in place, the biological interpretation becomes intuitive: the query ATAC-seq dataset from a tumor sample provides a “snapshot” of the genome-wide binding activities of functional TRs, akin to distinct ‘fingerprints’ at a crime scene, pointing toward individuals involved. The ChIP-seq database acts as reference library for TR identification, using stored datasets as distinctive "fingerprints" for individual TRs. By matching the tumor’s accessibility fingerprint against the reference library, eTRex identifies which TRs are most plausibly driving regulatory activity in that sample, much like matching evidence from a crime scene to known suspects in a database.

Second, it is noted many TRs have been profiled repeatedly, often due to the uneven allocation of research resources and attention. Some well-studied TRs (e.g., CTCF, TP53) can even be represented by hundreds of ChIP-seq datasets, whereas the majority have only one or two.

Performing independent enrichment analyses for each TR ChIP-seq dataset within accessible chromatin regions can therefore lead to overfitting and biased inference, particularly for TRs with more datasets. eTRex thus employs a Bayesian hierarchical probit framework (**Figure 1b**) that integrates consistency measures across these datasets to infer an overall TR-level consistency score. This hierarchical structure also enables information sharing among TRs, thereby stabilizing estimation and mitigating overfitting.

Next, eTRex employs a coordinate ascent algorithm to estimate the model parameters by minimizing the Kullback-Leibler (KL) divergence between the mean-field variational family and the true, complex posterior distribution^29^ (**Figure 1c**). Unlike BIT, which is based on a two-level logit model for count data, eTRex introduces a three-level probit model for binary data (**Figure 1b**), under which our coordinate ascent variational inference (CAVI) yields closed-form equations for updating all component variational distributions, offering both statistical rigor and computational efficiency. Finally, eTRex generates a functional TR profile for each input dataset by ranking TRs according to their inferred genome-wide consistency. Using this methodology, we generated functional TR profiles for 4,819 cancer datasets and established an online portal designed to offer user-friendly access to eTRex’s data resources and core analytical capabilities (**Figure 1d**).

A key advantage of this variational Bayesian approach is its substantially higher computational efficiency compared to stochastic sampling methods. By leveraging this advantage, eTRex achieves a drastic improvement in processing speed while maintaining comparable performance in identifying TRs. As a result, eTRex enables a comprehensive survey of the functional TRs across 4,819 datasets spanning 19 cancer types. The scalability of the eTRex algorithm makes it feasible to conduct the reported pan-cancer analysis task at unprecedented breadth, providing a solid foundation for uncovering shared and cancer-specific transcriptional regulatory programs.

To validate the proposed model, we first benchmarked eTRex against BIT and five other available methods capable of identifying TRs from accessible chromatin regions, including ChIP-Atlas^13^, WhichTF^17^, BART^18^, i-cisTarget^19^, and HOMER^20^, We used differentially accessible regions (DARs) derived from eight publicly available TR-perturbation experiments—six from human and two from mouse^37–41^ (**Figure 2a**). For mouse experiments we used a different reference library that contains 5,681 TR ChIP-seq datasets covering 607 mouse TRs, also sourced from GTRD. These DARs represent genomic regions showing altered chromatin accessibility after perturbation of a TR, and these perturbed TRs thus serve as ground truth for evaluating method performance to identify functional TRs.

**Figure 2.**
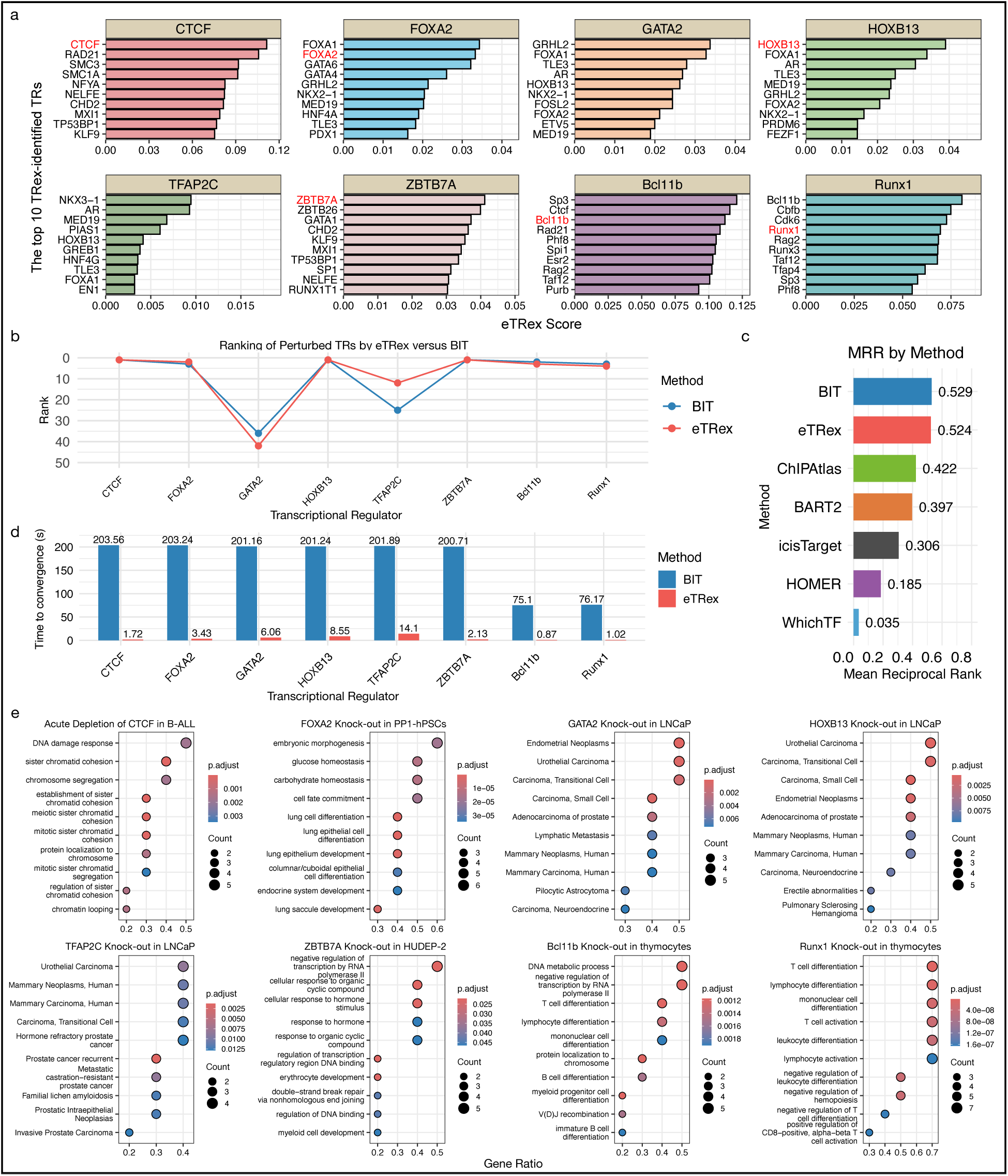
Benchmarking eTRex using differentially accessible regions (DARs) from TR perturbation experiments. (a) Top 10 TRs identified by eTRex across eight DAR datasets from TR perturbation experiments. (b) Comparison of perturbed TR rankings between eTRex (variational inference) and its predecessor BIT (MCMC). (c) Mean reciprocal rank (MRR) for perturbed TRs across eight ranking lists, calculated for eTRex, BIT, ChIP-Atlas, BART2, I-cisTarget, HOMER, and WhichTF. (d) Comparison of time to convergence between eTRex and BIT. (e) Gene ontology (GO) enrichment analysis of the top 10 TRs identified by eTRex, demonstrating strong enrichment in context-specific biological processes.

We first benchmarked the proposed eTRex method with the previously published BIT method, which shows these two Bayesian methods produced nearly identical rankings for six of the eight perturbed TRs (**Figure 2b**), despite their distinct model structure and setup. We then compared eTRex and BIT with the other five computational methods and summarized the results using mean reciprocal rank^13,17–20^ (MRR), a metric measuring the ability to identify the perturbed TR (**Figure 2c**). eTRex outperformed the five methods while achieving superior MRR performance comparable to BIT. Furthermore, a direct comparison of convergence times shows that eTRex is dramatically faster than BIT, achieving a speed-up factor ranging from approximately ×14 (for TFAP2C) to over ×118 (for CTCF) with an average speed-up of ×63 (**Figure 2d** and **Supplementary Figure 2**). We further show that in terms of the total runtime (including data preprocessing time), eTRex and BART are clearly the fastest, typically completing the analysis for each set of DARs in 0.7-1.5 minutes (**Supplementary Figure 3**). In contrast, the next fastest method, BIT, took 3.0-7.9 minutes, while the other methods could require well over one hour. eTRex thus achieves significantly higher computational efficiency while maintaining the top-level predictive accuracy.

eTRex can not only perform well in identifying perturbed TRs (**Figure 2a, 2d**) but also uncover previously unrecognized genes and regulatory pathways involved in the same process. For instance, for known prostate-cancer-specific TRs (e.g., GATA2, HOXB13, and TFAP2C)^42–44^, eTRex can leverage ATAC-seq data from these TR knockout experiments to identify a set of top-ranked TRs, which were then proven to be significantly enriched in prostate cancer–related gene ontology (GO) terms (**Figure 2e**), further validating eTRex’s ability to robustly identify functionally relevant TRs. As a result, it demonstrates eTRex’s potential in offering new insights for functional studies from analyzing related biological experimental data.

### eTRex-identified pan-cancer transcriptional regulators

To define a pan-cancer atlas of functional TR profiles, we collected and pre-processed 4,819 publicly available ATAC-seq datasets spanning 19 different cancer types (**Supplementary Figure 4**), sourced from ChIP-Atlas^13^. Previous studies that analyzed chromatin accessibility of various cancers have shown that active regulatory elements are highly enriched with TR binding sites that are linked to genes critical for cancer and tissue identity^24,45^. Guided by this insight, we focused our analysis on accessible chromatin regions overlapping ENCODE-annotated cis-regulatory elements^15^, thereby enriching for functionally relevant loci and improving the efficiency and precision of identifying functional TRs. This large-scale integration establishes a unique foundation for constructing a comprehensive atlas of oncogenic regulatory activity across human cancers.

We applied eTRex to these pan-cancer ATAC-seq datasets to generate a pan-cancer atlas aimed at revealing oncogenic transcriptional regulatory programs. To obtain a pan-cancer overview, we first classified the eTRex-derived 4,819 TR profiles by the 19 cancer types. A key analytical challenge lies in summarizing these TR profiles across diverse biological contexts. For example, the atlas contains 781 TR profiles derived from ATAC-seq datasets associated with breast cancer. To address this, we employed a flexible rank aggregation strategy. This method uses a mean reciprocal rank fusion (MRRF) to combine the rankings from all individual TR profiles within a given group (see **Methods**). We chose this rank-based approach because it is more robust to outliers and score variations than the alternative of directly averaging eTRex scores. Consequently, a high MRRF value indicates that a TR is consistently prioritized as functionally important across datasets within the same cancer type—offering a principled way to define consensus regulatory programs.

Based on the calculated MRRF for each TR within every cancer type, we stratified TRs into three confidence categories: high-confidence (MRRF > 0.01), corresponding to TRs that consistently rank in the top 50; medium-confidence (0.005 < MRRF ≤ 0.01), for TRs frequently ranking in the top 100 or occasionally ranked in the top 50; and low-confidence (MRRF ≤ 0.005), for those rarely prioritized by the eTRex method. The distributions of ranked positions for TRs in these three categories are illustrated by density plots (**Supplementary Figure 5**). The selected MRRF cutoffs are visually justified as they effectively segregate the TRs into distinct rank distributions: the high-confidence group shows a sharp peak concentrated primarily left of the top 50 rank line, the medium-confidence group peaks near the top 100 rank line, and the low-confidence group exhibits a nearly flat distribution across the entire rank range.

To further characterize the cross-cancer prevalence of high-confidence TRs, we examined their recurrence across cancer types—defining ubiquitous drivers if high-confidence in five or more cancer types (a threshold implicated by previous pan-cancer analyses^46^), intermediate-specific TRs when shared by two to four lineages, and highly specific TRs when restricted to a single cancer type (**Figure 3a**).

**Figure 3.**
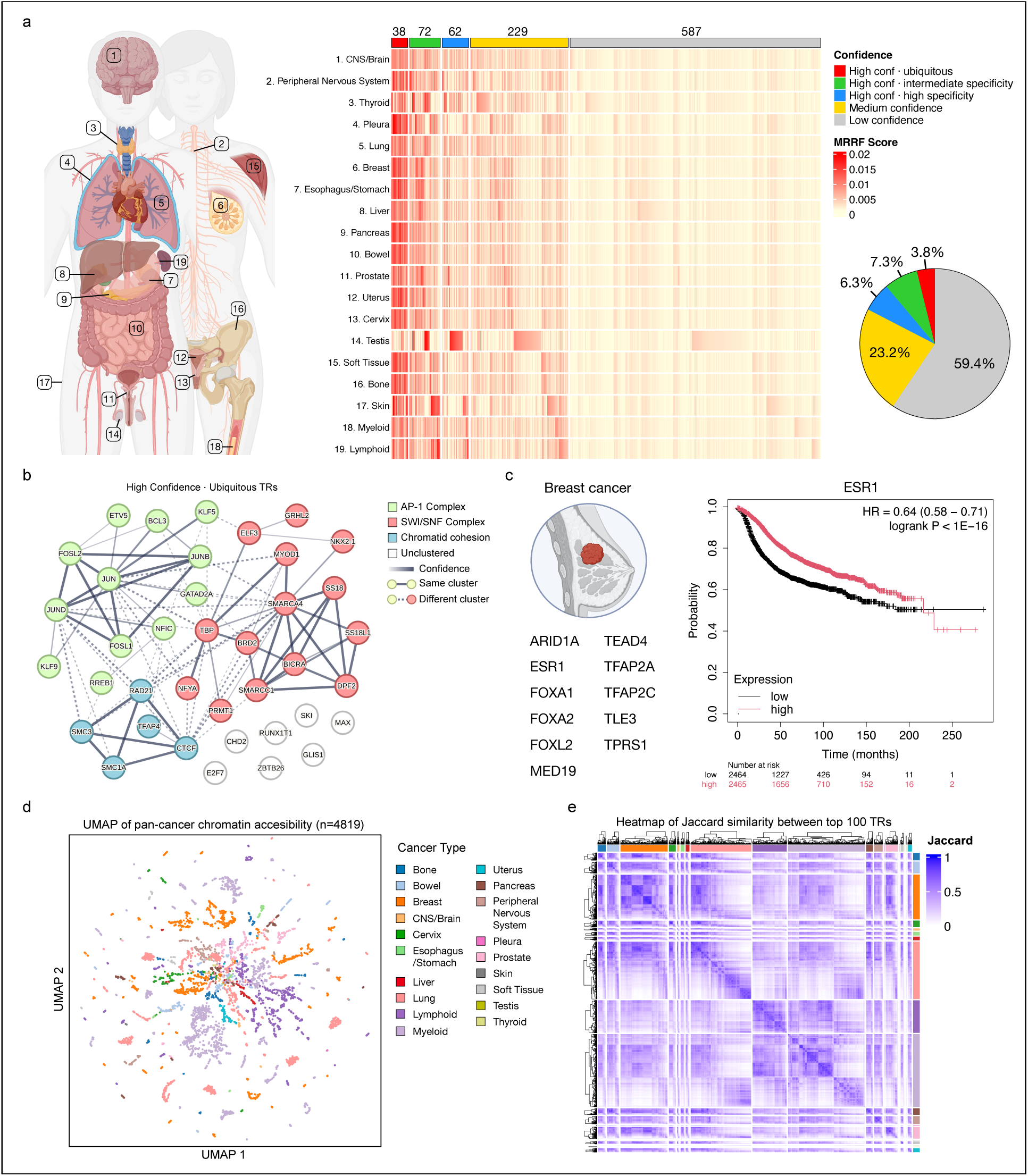
Pan-cancer functional transcriptional regulators identified by eTRex from chromatin accessibility data. (a). Pan-cancer functional transcriptional regulator (TR) profiles identified from chromatin accessibility data by eTRex through integration of cancer chromatin accessibility data and TR ChIP-seq data. TR profiles were grouped by cancer types and aggregated using mean reciprocal rank fusion (MRRF). (b). High-confidence (MRRF > 0.01) and ubiquitous TRs (MRRF > 0.01 in more than five cancer types), clustered using STRINGdb. (c). Cancer-type-specific TRs identified in breast cancer, and Kaplan Meier plot of ESR1 in breast cancer. (d). UMAP projection of 4,819 chromatin accessibility datasets, colored by cancer type. (e). Jaccard similarity heatmap computed from the top 100 TRs across the 4,819 TR profiles grouped by cancer types indicated by the color bars on the right and top sides of the heatmap.

We first examined ubiquitous high-confidence TRs, which clustered into three functional groups according to STRINGdb^47^: AP-1 complex, SWI/SNF chromatin-remodeling complex, and chromatid cohesion complex (**Figure 3b**). The consistently high eTRex rankings of these regulators across cancers underscore their core oncogenic relevance. For example, the AP-1 complex integrates diverse extracellular signals, orchestrating transcriptional programs that drive tumor proliferation, progression, and therapy resistance^48–50^. Similarly, the SWI/SNF complex controls nucleosome positioning and enhancer activity^51,52^, and mutations in its subunits occur in nearly 25% of all human cancers^53,54^.

Second, we focused on cancer-type-specific high-confidence TRs, which are likely to represent context-dependent oncogenic regulators. For example, in breast cancer, eTRex identified ARID1A, ESR1, and FOXA1 as top-ranked TRs. These are well-established drivers of estrogen receptor (ER)-positive tumorigenesis^55–57^. Specifically, ESR1 governs ER-positive breast cancer signaling^55^. FOXA1 acts as a pioneer factor to modulate chromatin accessibility for ESR1 binding^56^. ARID1A regulates genome-wide ER-FOXA1 chromatin interactions and ER-dependent transcription and has a critical role in maintaining luminal cell identity and endocrine therapeutic response in ER-positive breast cancer^57^. Moreover, a Kaplan–Meier analysis of >30,000 tumor samples showed that 8 of 11 eTRex-identified TRs were significantly associated with patient survival^58^ (**Figure 3c** for ESR1 and **Supplementary Figure 6** for all the 11 TRs), highlighting their potential as prognostic biomarkers. Nonetheless, many of the eTRex-identified cancer-type-specific TRs remain under-investigated, and further exploration of these TRs may provide valuable opportunities to discover and characterize novel oncogenic regulatory programs.

Third, to explore potential mechanistic links between regulatory disruption and tumorigenesis, we examined the overlap between the consensus binding sites of each TR and somatic mutation sites (sourced from DepMap^59^). Overall, high-confidence TRs exhibited significantly elevated mutation frequencies within their binding sites (**Supplementary Figure 7**), suggesting these regulatory regions are preferentially targeted by somatic alterations. Given that mutation accumulation in regulatory DNA is a known driver of cancer^60^, these findings implicate TR binding site perturbation as a potential mechanism of transcriptional dysregulation in oncogenesis.

Finally, the analysis of pan-cancer TR profiles aggregated by cancer type revealed both common and cancer-type-specific TRs. However, UMAP projection of binary vectors of ATAC-seq profiles also revealed pronounced heterogeneity within individual cancer types (**Figure 3d**), reflecting genetic and epigenetic divergence among subtypes. For example, luminal and basal breast cancers can display distinct accessibility and TR activity patterns^61^. Luminal subtypes depend on TRs such as ESR1 and FOXA1^61^, whereas basal subtypes are driven by AP-1 complex factors^62^. Consistently, pairwise Jaccard similarity of top ranked TRs revealed strong intra-type variation (**Figure 3e**). This underscores a major advantage of our resource: unlike existing studies that provide only aggregated, cancer-type-level summaries, the eTRex atlas delivers dataset-level context-specific TR profiles, enabling precise and customizable analyses at the cell line or subtype level or any user interested research tasks. By re-aggregating these profiles, researchers can flexibly investigate TR activity within any chosen biological context— offering a powerful, extensible resource for decoding oncogenic regulatory programs across human cancers.

### Aggregating functional TR profiles of leukemia cell lines

We now demonstrate how this inherent flexibility provides additional insights into context-specific oncogenic regulatory programs by aggregating TR profiles derived from same cancer cell line. Specifically, 4,819 functional TR profiles can be re-aggregated into 227 distinct cancer cell lines. To illustrate the utility of this approach, we compared two leukemia cell lines (K562 and Jurkat) with different cellular origins to highlight how our method can effectively uncover divergent oncogenic transcriptional regulatory programs.

Leukemia is a group of hematological cancers characterized by uncontrolled proliferation of abnormal blood cells, primarily affecting the bone marrow and circulating blood. The disease can be classified into acute and chronic forms, as well as myeloid and lymphoid cell lineages, depending on the lineage and maturity of the malignant cells involved. Two widely studied leukemia cell lines are K562 and Jurkat. K562 cells originate from a chronic myeloid leukemia (CML) patient and serve as a valuable model for studying erythroid differentiation and myeloid leukemia biology^63^. Jurkat cells, derived from an acute T-cell leukemia patient, are commonly used to investigate T-cell receptor signaling pathways and acute lymphoblastic leukemia (ALL) pathogenesis^64^.

Our analysis is based on TR profiles from 588 ATAC-seq datasets associated with K562 cells and 136 associated with Jurkat cells in the eTRex atlas. A UMAP projection generated from open chromatin regions confirmed a clear separation between these two cell lines (**Figure 4a**), reflecting their distinct chromatin accessibility landscapes and TR binding activities. We computed the MRRF for each TR in two cell lines, separately (**Figure 4b**). As anticipated, this analysis yielded distinct sets of high-confidence TRs (MRRF > 0.01). For instance, high-confidence TRs in K562 include CTCF, GATA1, KLF1, LDB1, MECOM, NFE2, SKI, ZBTB7A, and ZEB2, consistent with their established roles in erythroid development. In contrast, high-confidence TRs in Jurkat include well-characterized T-cell lineage-specific regulators such as ETS1, IKZF1, IRF4, MYB, RUNX1, SPI1, STAT5B, TBX21, TAL1, and TLX1, which are known regulators of lymphocyte development and function. The functional relevance of these TRs in leukemia is supported by previous studies (**Supplementary Table 1**). Beyond literature cross-validation, we conducted a disease enrichment analysis using the high-confidence TRs identified for each cell line. This analysis revealed significant enrichment in terms related to leukemia, corroborating the cellular origins. Notably, ’myeloid leukemia’ emerged as the primary term associated with eTRex-identified TRs from the K562 datasets, whereas ’lymphoid leukemia’ was the top term enriched with eTRex-identified TRs from the Jurkat datasets (**Figure 4c**). Furthermore, a gene concept network plot reveals the specific eTRex-identified TRs that mediate these leukemia-associated transcriptional regulatory programs^65^ (**Figure 4d**).

**Figure 4.**
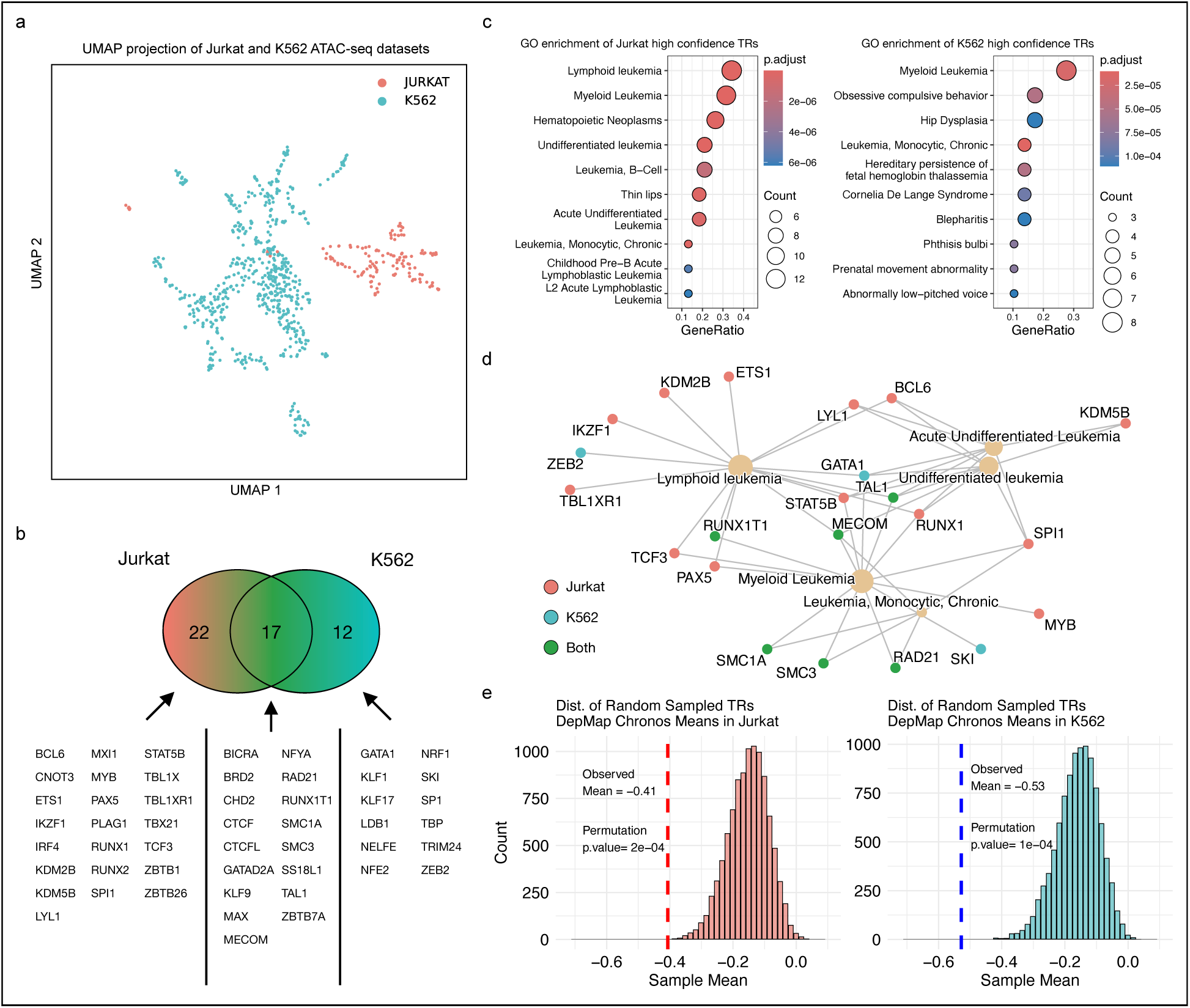
Contrasting TR profiles of K562 and Jurkat leukemia cell line. (a). UMAP projection of ATAC-seq datasets from Jurkat and K562 cell lines. (b). Venn diagram showing the overlap of high-confidence TRs between the two cell lines. (c). Gene Ontology (GO) enrichment analysis for high-confidence TRs identified in each cell line. (d). Gene concept network visualizing the association of eTRex-identified TRs with leukemia-related GO terms. (e). Permutation test results for the mean Chronos dependency scores of high-confidence TRs, compared against randomly selected TR sets of the same size.

Next, we sought to validate the functional significance of the eTRex-identified TRs using genome-wide cellular dependency data (Chronos scores, derived from CRISPR/Cas9 knockout screens) across numerous cancer cell lines, sourced from DepMap experimental datasets^59^. We specifically examined whether the mean Chronos scores of the high-confidence TRs deviated significantly from random expectation by permutation tests. These tests provided empirical p-values quantifying the likelihood of observing such mean values, or values more extreme, under the null hypothesis of random ranking (**Figure 4e**). The results clearly showed that the observed mean Chronos score for these eTRex-identified TRs was significantly lower (indicating higher dependency) than random expectation (K562: 𝑝 < 0.001, Jurkat: 𝑝 < 0.001). This finding strongly suggests that the high-confidence TRs tend to be functionally essential in these two leukemia cell lines.

To extend this validation, we used MRRF to aggregate eTRex-defined TR profiles for all 227 distinct cancer cell lines derived from the 4,819 cancer datasets. For the 155 cancer cell lines with available Chronos scores in DepMap, we repeated the permutation test to assess the biological relevance of the high-confidence TRs. After applying the Benjamini-Hochberg correction for multiple testing, 121/155 (78.1%) cell lines have adjusted p-value < 0.05 (**Supplementary Figure 8**). Collectively, this extended validation analysis across hundreds of diverse cell lines demonstrates that eTRex-prioritized TRs are consistently and significantly associated with increased cellular dependency, strongly supporting their functional importance in various cancers.

### Aggregating functional TR profiles of breast cancer subtypes

In addition to cancer cell lines, TR profiles in the atlas can also help characterize transcriptional regulatory programs across distinct cancer subtypes. Here, we reaggregated 781 breast-cancer-associated TR profiles into four major clinically relevant subtypes^61^: normal-like (13.7%), luminal (55.2%), HER2-enriched (10.2%), and basal-like (20.8%). The UMAP projection based on chromatin accessibility revealed clear distinctions among these four subtypes (**Figure 5a**), implying differences in their underlying TR activities.

**Figure 5.**
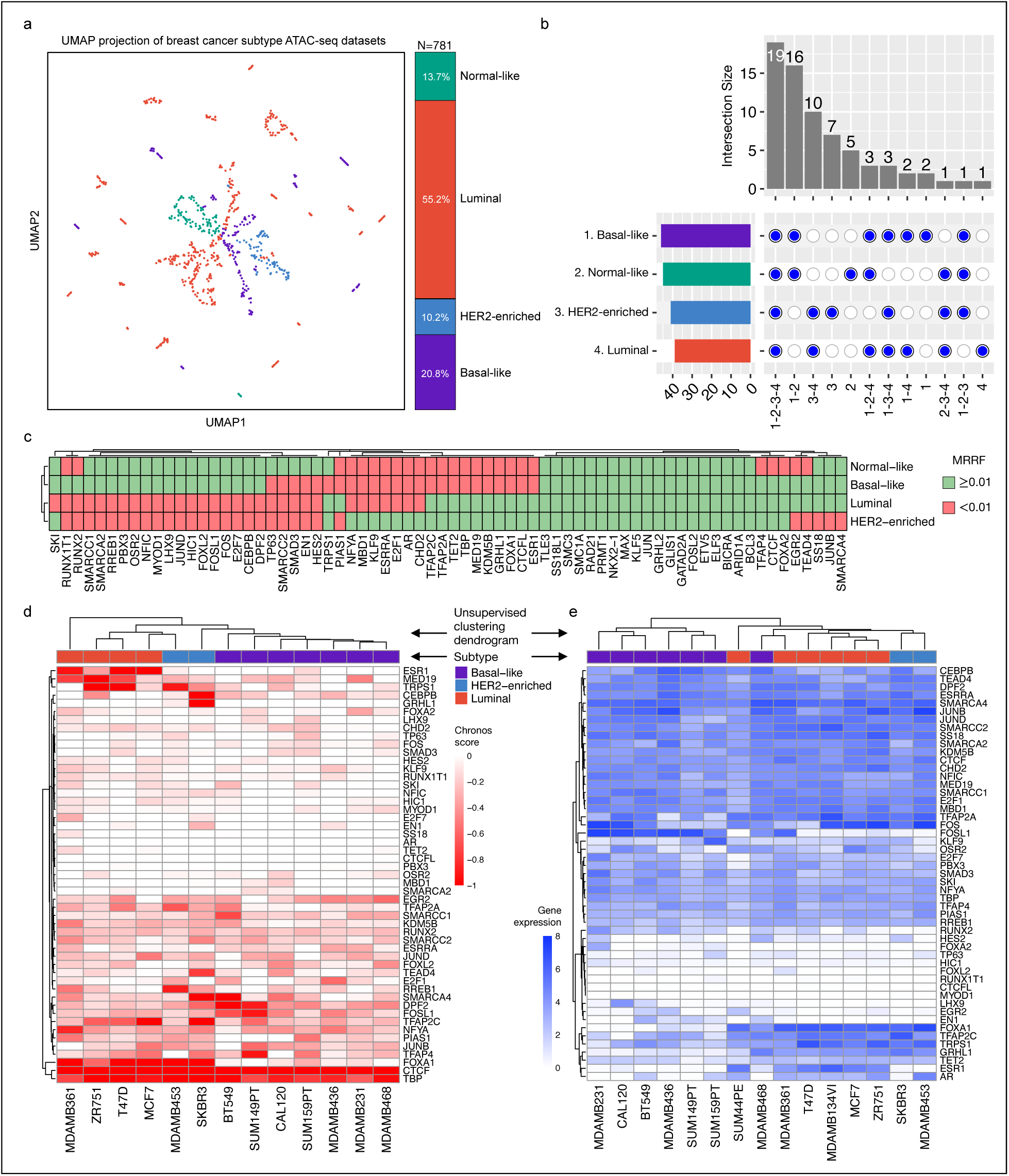
eTRex-identified transcriptional regulators across breast cancer subtypes. (a) UMAP projection of breast cancer ATAC-seq datasets, with points colored by subtype. (b) UpSet plot displaying the overlap of high-confidence TRs among the four subtypes. (c) Mean reciprocal rank fusion (MRRF) scores for high-confidence TRs across the four breast cancer subtypes. (d) Unsupervised hierarchical clustering of 13 breast cancer cell lines using their Chronos dependency scores for subtype-specific high-confidence TRs. (e) Unsupervised hierarchical clustering of 15 breast cancer cell lines using their gene expression levels for subtype-specific high-confidence TRs.

We first computed the MRRF scores separately for each TR within the four breast cancer subtypes and compared the high-confidence TRs (MRRF > 0.01). Our analysis indicated that, besides a core set of 19 common regulators shared by all four subtypes, luminal and HER2-enriched subtypes shared 10 high-confidence TRs, whereas the basal-like subtype shares 16 high-confidence TRs with normal-like subtype (**Figure 5b**). This finding aligns with established knowledge: luminal and HER2-enriched subtypes share a common origin from luminal epithelial cells of the mammary gland, while basal-like tumors often display transcriptional and regulatory features associated with progenitor-like or less differentiated epithelial states^66^. The high overlap of high-confidence TRs between basal-like subtype and normal-like subtype is also consistent, since the normal-like category mainly consists of TR profiles generated from the MCF10A cell line, which exhibits a basal-like phenotype^67,68^.

Next, we examined the high-confidence TRs within each breast cancer subtype and observed that, beyond common regulators such as the chromatin cohesion proteins and SWI/SWF complex proteins, subtype-specific high-confidence TRs include well-established breast cancer subtype markers. For instance, ESR1, FOXA1, and TET2 were identified by eTRex as high-confidence TRs within the Luminal and HER2-enriched subtypes^69–71^, while absent from basal-like and normal-like groups. eTRex also highlighted E2F1, GRHL2, TFAP2A, and TFAP2C, which are well-known regulators crucial to the HER2-enriched subtype^72–74^.

In contrast, the tumor suppressor TP63 was classified as high-confidence specifically within normal-like cells but not in the other subtypes, consistent with its role in maintaining epithelial identity and its high expression in MCF10A-derived non-malignant models^75^. The basal-like subtype was characterized by AP-1 complex TRs (**Figure 5c**) that promote aggressive phenotypes like proliferation and invasion. Studies show that components of the AP-1 complex, such as FOS, FOSL1, JUNB, and JUND, have been implicated in basal-like breast cancer, where elevated AP-1 activity is linked to aggressive behavior and poor prognosis^76–78^. We also note that many other high-confidence TRs identified by eTRex remain under-investigated, requiring further characterization of their roles within specific breast cancer subtypes.

Finally, we again cross-validated the cancer-subtype-specific high-confidence TRs with the DepMap experimental data. The 781 eTRex-defined TR profiles can be classified into 16 breast cancer cell lines, 13 of which have corresponding Chronos dependency scores in DepMap database. After excluding high-confidence TRs shared by all four subtypes, we performed unsupervised hierarchical clustering of these 13 cell lines using the remaining TRs’ dependency scores. Basal-like and luminal cell lines form clearly distinct branches, and the two HER2-enriched lines are also placed adjacently, consistent with their known subtypes (**Figure 5d**). In addition, applying the same approach to the 15 cell lines with corresponding TR gene expression levels in DepMap yielded a similarly robust subtype structure, with a nearly perfect separation of luminal and basal-like cell lines and again, the two HER2-enriched lines aligned together (**Figure 5e**). Together, these results indicate that the high-confidence TRs identified by eTRex capture the key features of breast cancer subtype identity and are biologically meaningful across both cellular dependency and gene expression data.

## Discussion

Understanding the heterogeneity of transcriptional regulation across cancers requires approaches that preserve context-specific regulatory information. To this end, we developed eTRex, a scalable variational Bayesian framework that integrates ATAC-seq–derived chromatin accessibility with experimentally measured transcriptional regulator (TR) binding sites to infer functional regulators. Applying eTRex to 4,819 datasets spanning 19 cancer types, we constructed a comprehensive pan-cancer atlas of functional TR profiles that meticulously maintains the distinct regulatory programs of each dataset. Unlike conventional methods that average or merge chromatin signals, the eTRex-derived atlas enables systematic exploration of both shared oncogenic programs and context-specific regulators defining individual cancer types, subtypes, cell lines, or any user-interested research tasks. This atlas represents a biologically interpretable, data-driven resource for dissecting the transcriptional regulatory landscape of human cancers.

eTRex introduces key advances in accuracy, scalability, and analytical flexibility in inferring functional TRs. It complements sequence-based, motif-based, and expression-based TR inference strategies by directly incorporating experimentally measured binding information, thereby improving discrimination among closely related regulators and offering an interpretable alternative to complex deep learning models. It relies on a novel three-level hierarchical probit model that accounts for between- and within-TR heterogeneity and mitigates bias and overfitting. Because eTRex infers posterior distributions over all model parameters, it also offers uncertainty quantification, allowing users to assess the robustness of regulatory scores in individual datasets.

Unlike MCMC-based approaches that require iterative sampling, our variational Bayesian formulation admits closed-form parameter updates, achieving both analytical simplicity and substantial computational gains. Replacing computationally intensive MCMC sampling with variational optimization enables efficient, large-scale inference across thousands of epigenomic datasets. As Bayesian methods continue to gain prominence in modern computational research^79,80^, the issue of slow convergence in traditional sampling-based algorithms deserves heightened attention, especially given the exponential growth of data.

Although this study focuses on constructing and applying a pan-cancer atlas, the eTRex framework is broadly applicable to any biological system where ATAC-seq or similar open chromatin data are available, including studies of development, differentiation, and disease. Because eTRex provides dataset-level context-specific TR profiles that can be flexibly re-aggregated, researchers can tailor analyses to any biological question, making it a versatile and extensible tool to create any new atlas.

Despite these advances, it is important to acknowledge the inherent limitations of all computational approaches, including eTRex. While such methods provide direct mechanistic insights, they cannot fully capture the complexity of transcriptional regulation, which is shaped by cooperative binding events, cofactor interactions, and chromatin states^81,82^. Consequently, even gold-standard ChIP-seq experiments provide only a snapshot of a TR’s potential activity. Accurately identifying the complete set of functional TRs under any given biological context therefore presents a significant challenge for the field. Nevertheless, by leveraging multiple ChIP-seq datasets for each TR and explicitly modeling both within- and between-TR heterogeneity, eTRex helps mitigate the challenge.

A second consideration involves aggregation across datasets. While we mainly used mean reciprocal rank fusion (MRRF) as the metric to aggregate TR profiles for user-defined biological contexts, other rank aggregation strategies can also be applied. The choice of aggregation method may influence the sensitivity to outliers or the weighting of individual datasets. Indeed, the problem of effectively combining multiple ranked lists has long been an active area of research in information retrieval and bioinformatics^83,84^. Future work may explore advanced rank aggregation approaches to further refine the integration of eTRex-derived results across datasets and contexts.

Another limitation relates to the potential influence of technical variation on context-specific features. While eTRex emphasizes the preservation of dataset-level chromatin accessibility features, some of the identified accessible chromatin regions may partially reflect technical or experimental biases rather than true biological features^35^. Such bias could arise from variations in sequencing depth, library preparation, or platform-specific effects^85^. Although proper preprocessing and quality-control steps can mitigate these issues, further refinement and harmonization of raw ATAC-seq data will be important for reducing technical confounding in subsequent analyses.

Looking forward, rapid advances in single-cell sequencing underscore the need to characterize transcriptional regulatory programs at unprecedented resolution. Single-cell ATAC-seq, often performed directly on heterogeneous tumor samples rather than cultured cell lines, captures the epigenetic heterogeneity inherent to malignant tissues^86^. Although most existing analyses focus on pseudo-bulk (cluster-level) transcriptional regulatory programs, recent studies reveal substantial heterogeneity even among cells within the same cluster^87^. Therefore, extending a scalable framework such as eTRex to single-cell settings represents a promising direction for future development, enabling functional TR profiling at both the cluster and per-cell levels.

Future integration with additional regulatory layers, such as histone modifications, multiome measurements, or CRISPR-based perturbation data, may further enhance the resolution and biological interpretability of eTRex-derived regulatory maps.

## Method

### Data Preprocessing

#### Analysis of Differentially Accessible Regions (DARs) from Bulk ATAC-seq

To identify differentially accessible chromatin regions from bulk ATAC-seq data in TR perturbation experiments, we employed the following computational pipeline. First, raw paired-end reads underwent quality assessment with FastQC (v0.12.1) and adapter trimming using Trim-galore (v0.6.10). The cleaned reads were then mapped to the appropriate reference genome (hg38 for human, mm10 for mouse) using bowtie2 (v2.5.2) with high-sensitivity parameters (--very-sensitive -X 2000). Post-alignment processing was conducted with SAMtools (v1.16.1) to convert SAM to BAM format, remove unmapped and duplicate reads, and create index files.

Subsequently, chromatin accessibility peaks were called from these processed alignments using MACS2 (v2.2.9.1). Finally, the resulting peak files and BAM files were used as input for the DiffBind R package (v3.10.1) to perform statistical analysis and identify significant DARs.

### Converting accessible chromatin regions and TR binding sites to binary vectors

For each TR perturbation dataset, we kept all differentially accessible regions generated by DiffBind. For each cancer ATAC-seq dataset, we first used “bedtools2 (v2.30.0)” to retain only regions overlapping with regulatory elements annotated by the ENCODE SCREEN project^15^. Next, the processed ATAC-seq dataset was provided as input to eTRex. All accessible chromatin regions from the input, as well as TR binding sites from each reference ChIP-seq dataset, were mapped to predefined, non-overlapping 1000 base-pair bins on the genome. Each bin was assigned either “0” or “1” according to whether the summit of any of the peaks falls into the bin (if the summit is missing, the middle point is used), transforming all datasets (input and reference) into the binary vectors of the same length. We then compared the binary vector from the input with that from each TR ChIP-seq dataset. For any given bin, we defined three conditions: (i) “matching” if both are (1,1); (ii) “mismatching” if they are (1,0) or (0,1); and (iii) “non-informative” if both are (0,0). In all input vs. reference comparisons, the first two conditions (“informative” bins) are much fewer than the third condition (“non-informative” bins). To address this sparsity while retaining valuable information, we dropped the “non-informative” bins.

### eTRex model and data augmentation method

eTRex is a Bayesian hierarchical probit model that leverages the strategy of data augmentation and implements variational inference to improve computational efficiency. We index TRs with multiple reference ChIP-seq datasets by 𝑖 = 1,2, … , ℳ, each having 𝑗 = 1,2, … , 𝐽_*i*_ datasets. We further index each informative bin when comparing the input to the 𝑗^th^ reference dataset of the 𝑖^th^ TR by 𝑘 = 1,2, . . , 𝑛_𝑖j_, where 𝑛_𝑖j_ denotes the number of informative bins. Similarly, TRs with a single reference dataset are indexed by 𝑖′ = 1,2, . . . , ℳ^𝑐^, all with 𝑗 ≡ 1 dataset, and their informative bins are indexed by 𝑘 = 1,2, … , 𝑛_*i*_′_1_.

Let 𝑥_𝑖j𝑘_ be the binary status of the 𝑘^th^ informative bin in the 𝑗^th^ reference dataset of the 𝑖^th^ TR, where 𝑥_𝑖j𝑘_ = 1 indicates “matching” and 𝑥_𝑖j𝑘_ = 0 indicates “mismatching”. The probability of an informative bin being “matching” is denoted by 𝑝_𝑖j_ (𝑝_*i*_′_1_), with 𝑃5𝑥_𝑖j𝑘_ = 1) = 𝑝_𝑖j_ (𝑃(𝑥_*i*_′_1𝑘_ = 1) = 𝑝_*i*_′_1_). Let *θ*_𝑖j_ = Φ^−1^(𝑝_𝑖j_) 5*θ*_*i*_′_1_ = Φ^−1^(𝑝_*i*_′_1_)) be the probit transformed 𝑝_𝑖j_(𝑝_*i*_′_1_), which is interpreted as the importance score measuring the consistency of binding patterns between input and reference at the dataset level. We further integrate information from multiple datasets of the same TR to get the importance score at the TR level (*θ*_*i*_/*θ*_*i*_′) by assuming 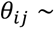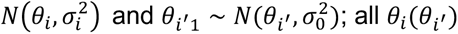; all *θ*_*i*_(*θ* ′) are governed by a global importance score 𝜇 and variance 𝜏^2^ by assuming *θ*_*i*_, *θ*_*i*_′ ∼ 𝑁(𝜇, 𝜏^2^). In this way, for each TR with multiple datasets, we model its dataset-level consistency scores using a normal distribution with distinctive mean *θ*_*i*_ and variance 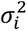, which reflect its overall consistency and within-TR heterogeneity, respectively. TRs with only one dataset share a common variance 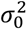 for information pooling.

The prior distributions over the hyperparameters follow standard choices in Bayesian hierarchical modeling: we place a weakly-informative prior on the global mean, that is 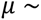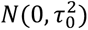, and assign gamma priors based on empirical knowledge learned from data to the precision terms 1/𝜏^2^ ∼ 𝐺𝑎𝑚𝑚𝑎(𝑎, 𝑏) and 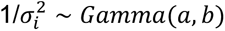 for 𝑖 = 0,1, … , ℳ. We specify the 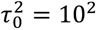, and 𝑎 = 𝑏 = 0.01 throughout this study. Under these specifications, the full hierarchical model can be written as:

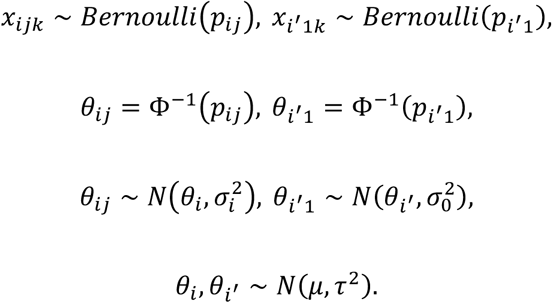

Notably, there is no conjugate prior exist for the probit transformed variable *θ*_𝑖j_. To overcome this problem, we leverage the data augmentation strategy by introducing a group of latent variables 𝑧_𝑖j𝑘_ and 𝑧_*i*_′_1𝑘_^88^. These latent variables follow Gaussian distribution centered at *θ*_𝑖j_ (*θ*_*i*_′_1_), and 𝑥_𝑖j𝑘_ (𝑥_*i*_′_1𝑘_) become deterministic conditional on the sign of 𝑧_𝑖j𝑘_ (𝑧_*i*_′_1𝑘_).

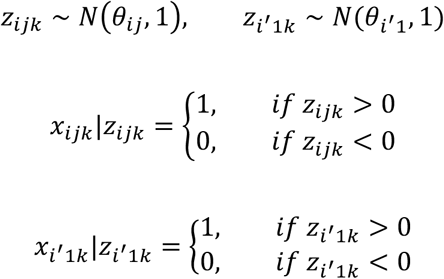

Let the observed data be 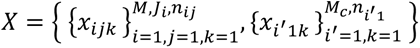. The unknown quantities, including parameters and latent variables involved in the eTRex model, are collectively denoted by 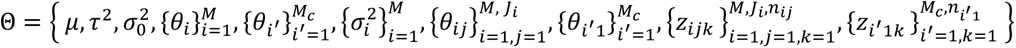.The full probability model of eTRex is then given by the product of all prior and conditional distributions:

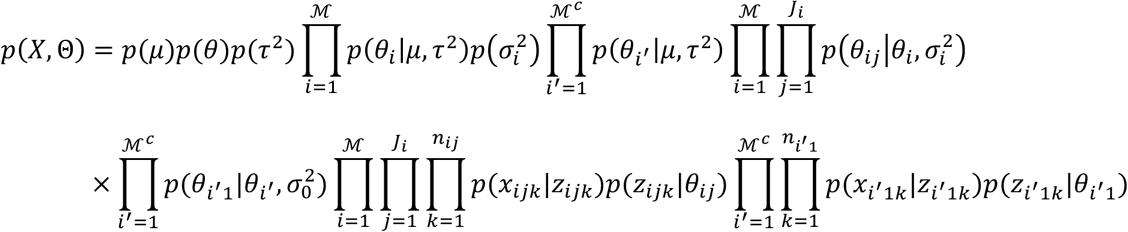

### Coordinate Ascent Mean-field Variational Inference

eTRex employs variational inference that recasts the problem of estimating the joint posterior distribution as an optimization problem: it seeks to minimize the KL divergence between the chosen variational distribution family and the true posterior distribution. eTRex uses a mean-field variational family which assumes that the latent variables and parameters are mutually independent, and each governed by a distinct factor in the variational distribution 𝑞(Θ),

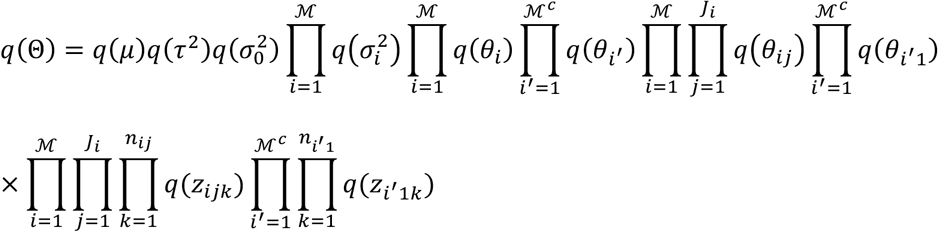

Within the mean-field variational family, CAVI iteratively updates each factor while holding all other factors fixed. The optimal solution of a factor has the general form,

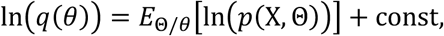

where the expectation is taken with respect to the current variational distributions of all remaining latent variables or parameters denoted by Θ/*θ*. For instance, based on the joint distribution 𝑝(𝑋, Θ) and factorized variational distribution 𝑞(Θ), the updates for 𝑞(*θ*_𝑖j_) and 𝑞(*θ*_*i*_′_1_) can be derived by applying this identity while keeping all other factors fixed:

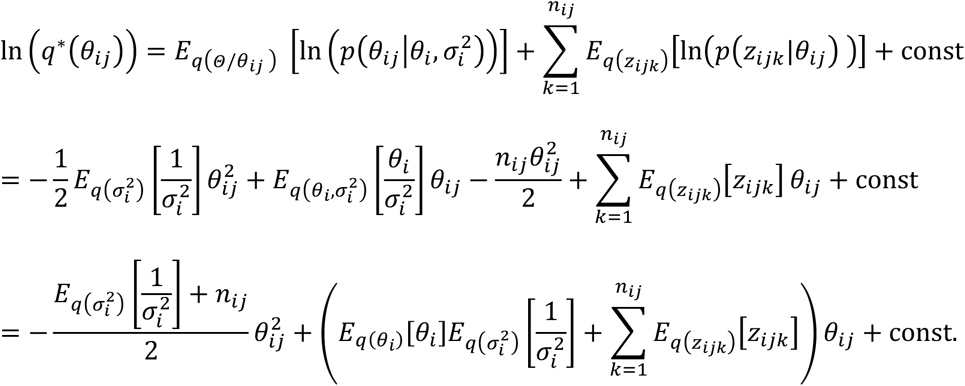

Therefore,

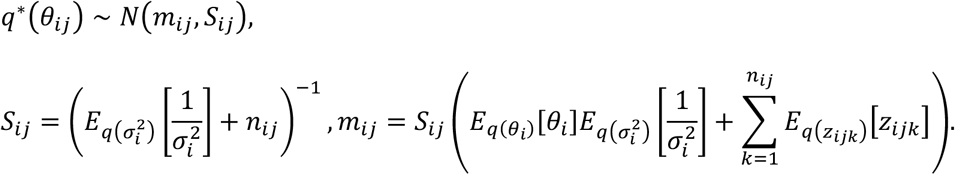

Similarly,

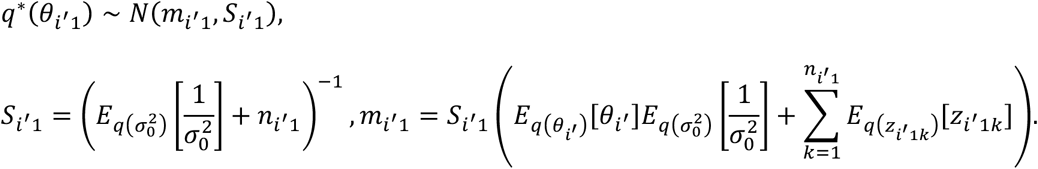

Optimal updates for the remaining factors can be derived in the same manner. To save the space, the complete derivations including all expectations, full coordinate-ascent update equations, and one-iteration update details are provided in **Supplementary Note 1**.

To robustly and flexibly aggregate the TR profiles, eTRex employs a rank-aggregation metric named “Mean Reciprocal Rank Fusion” (MRRF). MRRF is a modified version of the reciprocal rank fusion, specifically designed to enable cross-group comparisons of the same TR. Indexing each group as 𝑖 = 1, … , 𝐼 and within each group the TR profiles are indexed as 𝑗 = 1, … , 𝐼_j_, and ranking of the 𝑘^th^ TR is therefore denoted as 𝑅_𝑖j𝑘_. The MRRF of the TR in 𝑖^th^ group is calculated as

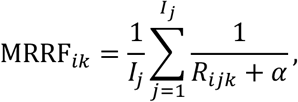

where 𝛼 is a smoothing factor (set to 50 throughout this study), analogous to the smoothing used in standard reciprocal-rank fusion. This measure can be interpreted as a normalized weighted Borda count, where reciprocal-rank weighting is applied and smoothed to prevent over-emphasizing the top regulators within a single profile. This smoothing factor balances contributions from different rank positions and prevents a single high-ranked TR profile from disproportionately influencing the results. Thus, MRRF provides a robust, balanced metric for aggregating TR rankings across heterogeneous groups.

### eTRex validation analysis

Benchmarking metric: To compare eTRex with existing methods, we use the mean reciprocal rank (MRR). Let 𝑅_*i*_ denote the rank assigned to the perturbed TR by a method in its 𝑖^th^ output list. For the eight perturbed TRs (CTCF, ZBTB7A, FOXA2, GATA2, HOXB13, TFAP2C, Runx1, and Bcl11b), the MRR of the method can be computed as:

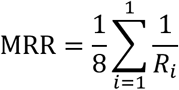

It is noted that MRR differs from the MRRF, as MRR is used as a metric to evaluates how effectively a method prioritizes the perturbed TR toward the top positions, while MRRF is used to aggregate multiple output ranking lists to assign a unified rank to each TR within a predefined category.

Evaluation of time to convergence: For BIT, convergence is evaluated using parameter trace plots and stabilization of hyperparameter 𝜇, as it governs all the TR-level parameters *θ*_*i*_ and *θ*_*i*_′. MCMC chains typically require roughly 500–1000 iterations of burn-in before reaching stationarity; therefore, we use 1000 iterations as the cutoff for defining convergence across all perturbation datasets. Similarly, convergence of the variational inference updates is monitored using parameter-difference trajectories, defined as Δ𝜇 = |𝜇^(𝑡)^ − 𝜇^(𝑡–1)^|, with a threshold of 0.001. In practical applications, we allow the variational updates to continue for a thousand iterations to ensure that the variational parameters remain stably converged.

Overlap with somatic mutation sites. To determine whether high-confidence TR binding sites are enriched for somatic mutations, we generate a consensus set of binding sites for each TR using bedtools2 (v2.30.0). For TRs represented by more than two ChIP-seq datasets, we retain a binding site in the consensus set only if it appears in at least two datasets. We then use the regioneR (v1.36.0) package in R to quantify, for each cancer type, the number of DepMap somatic mutation sites that fall within the consensus binding sites of each TR.

Cross-validation with DepMap experimental data: For each cancer cell line, we source CRISPR/Cas9 knockout screening data from DepMap, which measure cell viability after knocking out a specific TR gene and are summarized by the Chronos score (where lower scores indicate greater loss of viability). To evaluate whether high-confidence TRs are functionally essential, we perform permutation tests by randomly selecting 10,000 times of the same number of TRs as the eTRex-identified high-confidence TRs in each cell line and calculating the mean Chronos score for each random set. We then count how often the randomly selected TRs have a mean Chronos score lower than that observed for the high-confidence TRs. The proportion of such cases represents the empirical p-value, indicating how extreme the high-confidence TRs are relative to random expectation.

Gene ontology enrichment analysis. We use the R package clusterProfiler (v4.14.6) to perform gene ontology enrichment analysis. The top-ranked TRs in the BIT output list are first converted to their Ensembl IDs and then contrasted against the universe of all available genes to evaluate GO-term enrichment. We perform enrichment analysis for the top 10 TRs from the TR perturbation experiments and the high-confidence TRs from K562 and Jurkat. We apply a q-value cutoff of 0.05 and report the top 10 enriched terms in the results.

## Data Availability

Only public datasets were analyzed in this study. The pan-cancer ATAC-seq peak sets were retrieved from ChIP-Atlas (v3.0.0) [https://chip-atlas.org]. The human and mouse TR ChIP-seq peak sets were sourced from GTRD database (v21.12) [http://gtrd.biouml.org:8888/downloads/current/]. Cancer-type specific accessible regions were retrieved from TCGA^24^ (GDC 2019 v18.0, https://gdc.cancer.gov/about-data/publications/ATACseq-AWG). Raw ATAC-seq sequencing data from TR perturbation experiments are available at the GEO under accession codes: GSE153237^37^ [https://www.ncbi.nlm.nih.gov/geo/query/acc.cgi?acc=GSE153237], GSE173416^39^ [https://www.ncbi.nlm.nih.gov/geo/query/acc.cgi?acc=GSE173416], GSE114102^38^ [https://www.ncbi.nlm.nih.gov/geo/query/acc.cgi?acc=GSE114102], GSE234331^41^ [https://www.ncbi.nlm.nih.gov/geo/query/acc.cgi?acc=GSE234331], and GSE275927^40^ [https://www.ncbi.nlm.nih.gov/geo/query/acc.cgi?acc=GSE275927]. CRISPR/Cas9 knockout screening data, gene expression profiles, and somatic mutation data for each cell line were downloaded from DepMap [https://depmap.org/portal/].

## Code Availability

eTRex software is available at GitHub [https://github.com/ZeyuL01/eTRex] with a MIT license. We provided a web portal for online analysis [https://e-trex.org].

## Author Contributions

Lu, Z. conducted data-processing, coding, simulation, and validation. Lu, Z. and Wang, X. developed eTRex’s model and methodology and designed the algorithm and numerical studies. Wang, X. and Xu, L. conceived the ideas and supervised the study. Wang, X. acquired the funding. Yang, Y., Zheng, Q. and Gao, F. provided feedback on method or software development. All members have read, revised, and approved the final manuscript.

## Competing Interests

The authors declare no competing interests.

## Notes

### Competing Interest Statement

The authors have declared no competing interest.

