## Supplementary for "eTRex Reveals Oncogenic Transcriptional Regulatory Programs Across Human Cancers"

### eTRex Reveals Oncogenic Transcriptional Regulation Program Across Human Cancer

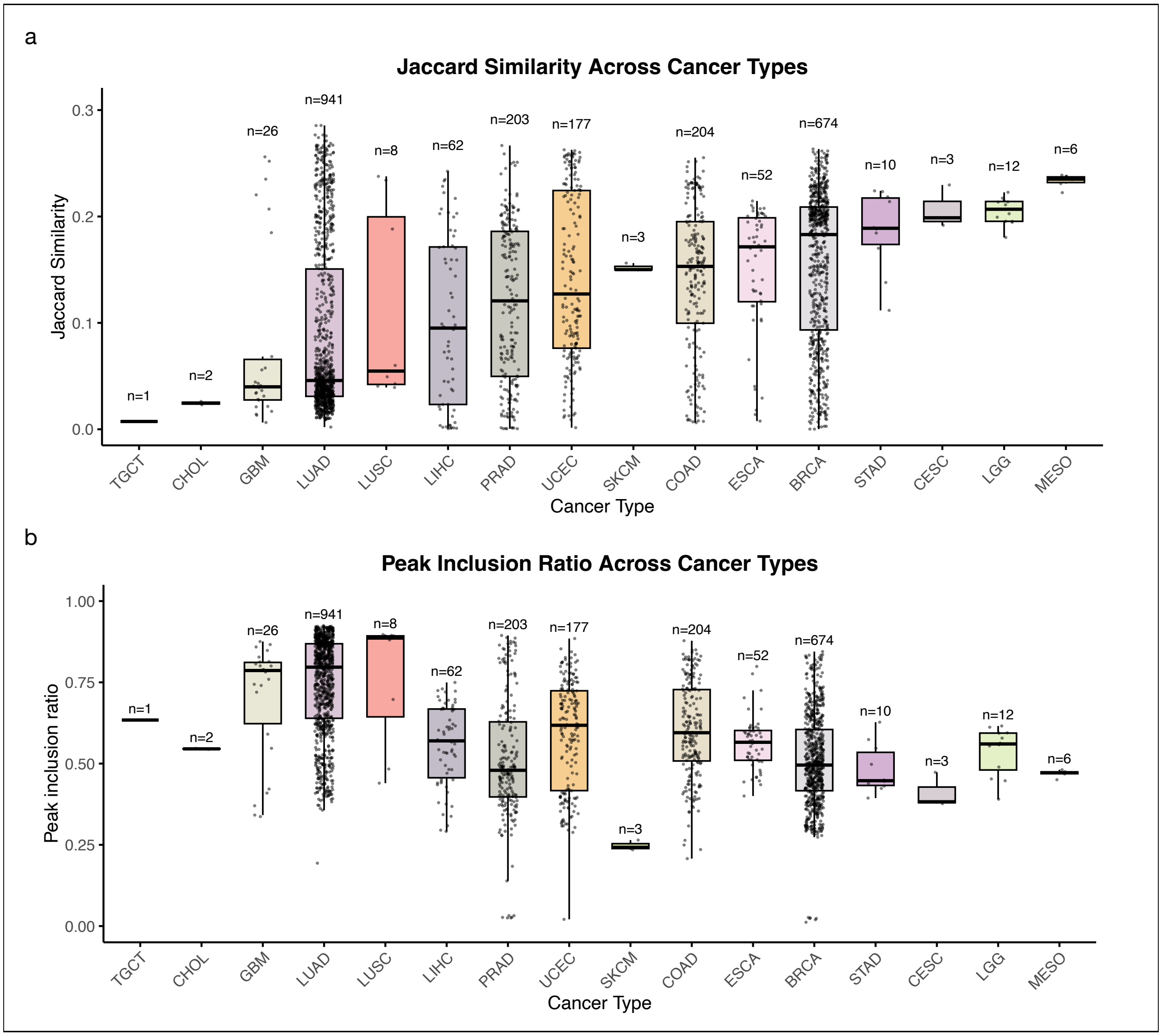


**Supplementary Figure 1. Variability in chromatin accessibility between individual ATAC-seq datasets and aggregated cancer-type–specific profiles.** (a) Jaccard similarity between chromatin accessible regions identified in each individual ATAC-seq dataset and the aggregated cancer-type–specific chromatin accessibility profiles reported in previous studies. (b) Proportion of peaks from each individual ATAC-seq dataset that overlap with the aggregated cancer-type–specific chromatin accessibility regions, reflecting the inclusion ratio of context-specific peaks within the aggregated profiles.

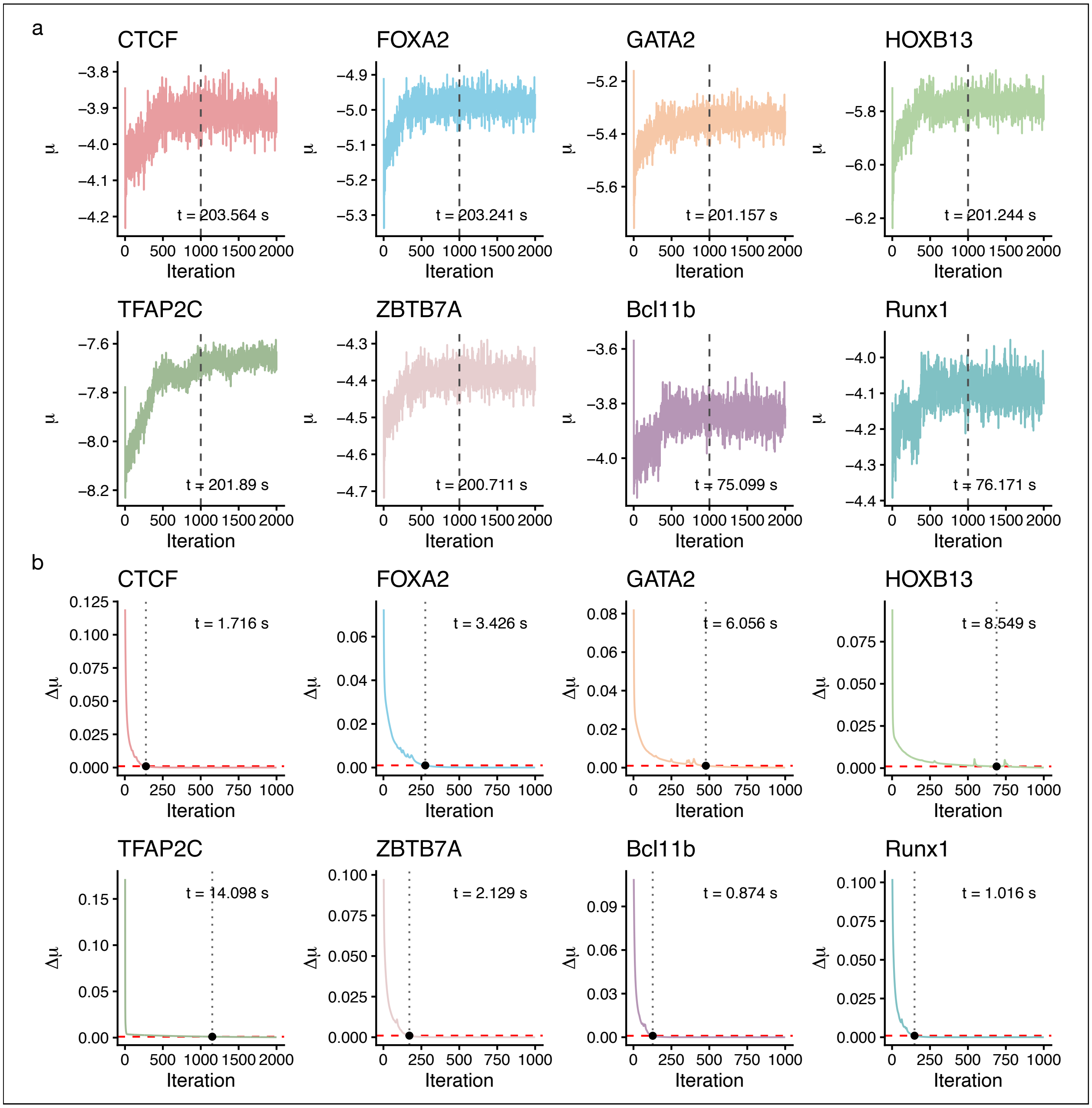


**Supplementary Figure 2. Convergence behaviors of BIT and eTRex on eight TR-perturbation experiments.** (a). BIT trace plots ($\mu_0$) showing stabilization after approximately 500–1000 burn-in iterations. (b). eTRex convergence was assessed using parameter-difference trajectories of the mean importance score $\mu$, with a change threshold of 0.001.

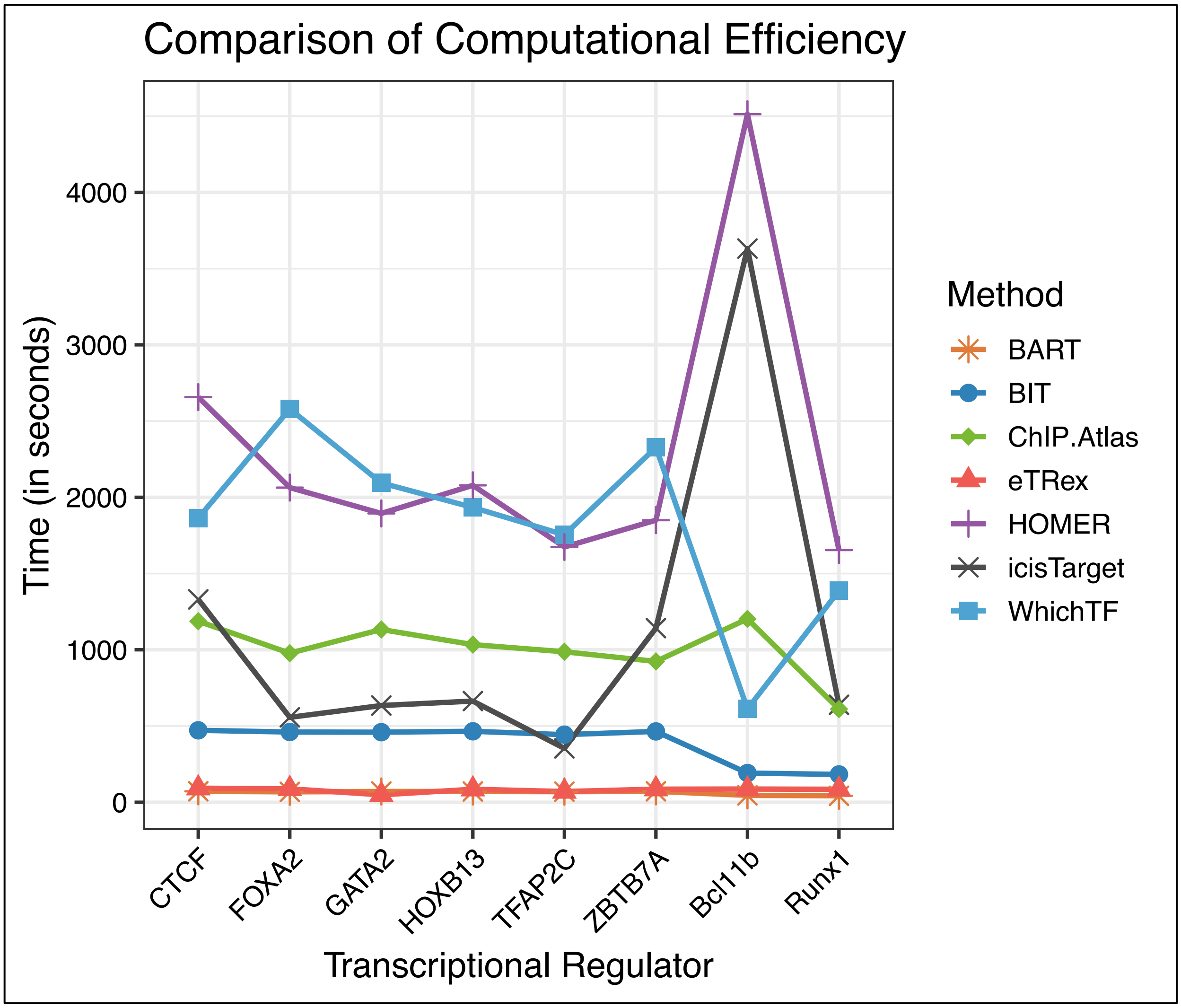


**Supplementary Figure 3. Comparison of computational efficiency**. Runtime comparison for eTRex, BIT, BART2, ChIP-Atlas, HOMER, i-cisTarget, and WhichTF across the eight datasets.

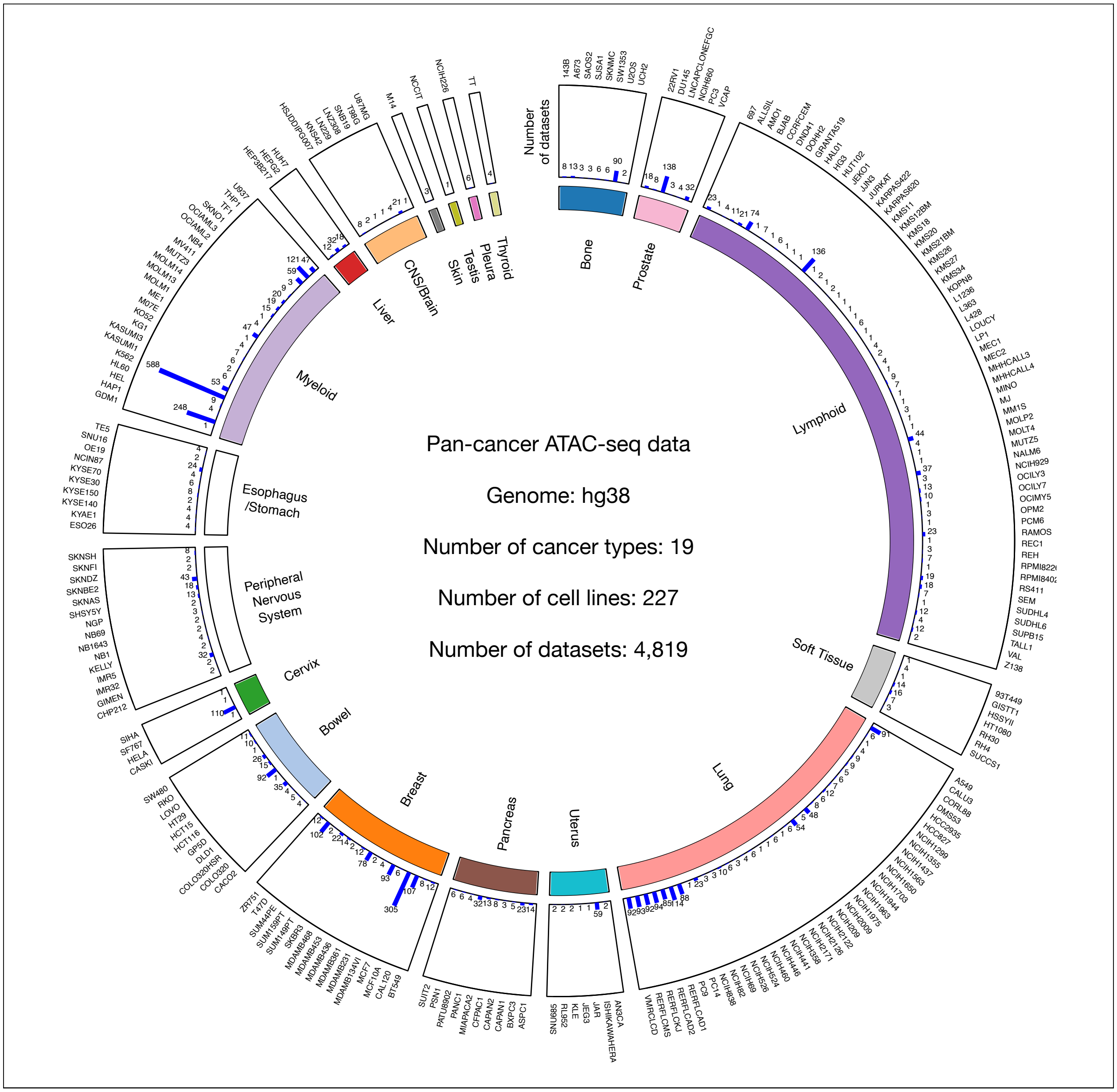


**Supplementary Figure 4. Overview of pan-cancer ATAC-seq data.** To identify pan-cancer TR profiles, we sourced 4,819 cancer-related ATAC-seq datasets from ChIP-Atlas, which can be associated to 19 cancer types or 227 different cancer cell lines.

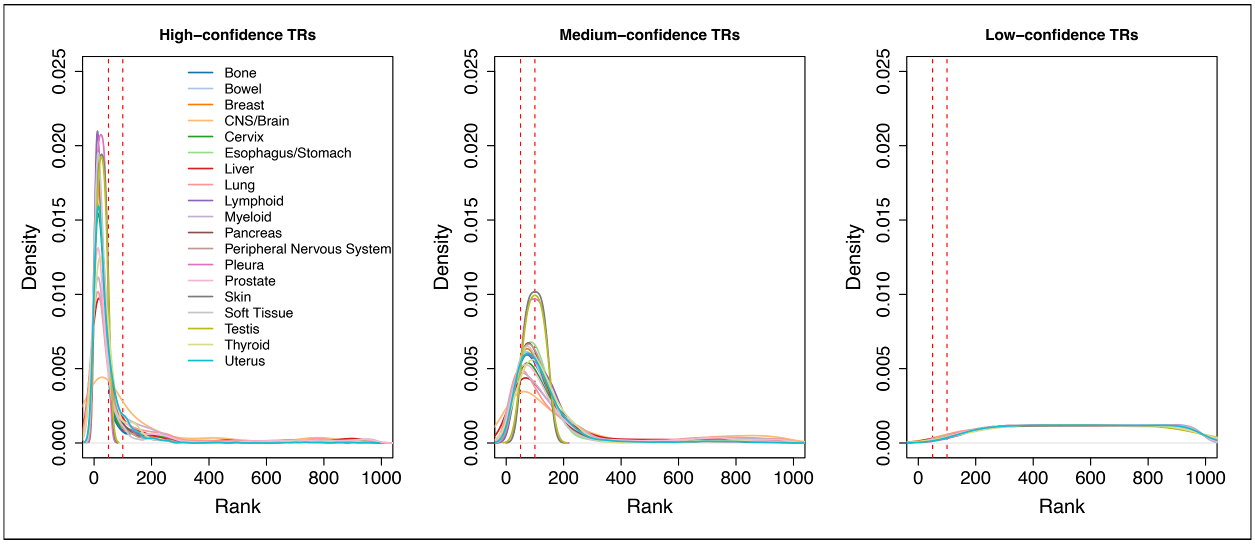


**Supplementary Figure 5.** Density plots of ranked positions for high-, medium-, and low-confidence transcriptional regulators (TRs), obtained by pooling the rankings from all TR profiles within each cancer type. The two dashed vertical lines indicate the cutoffs for the top 50 and top 100 ranked TRs.

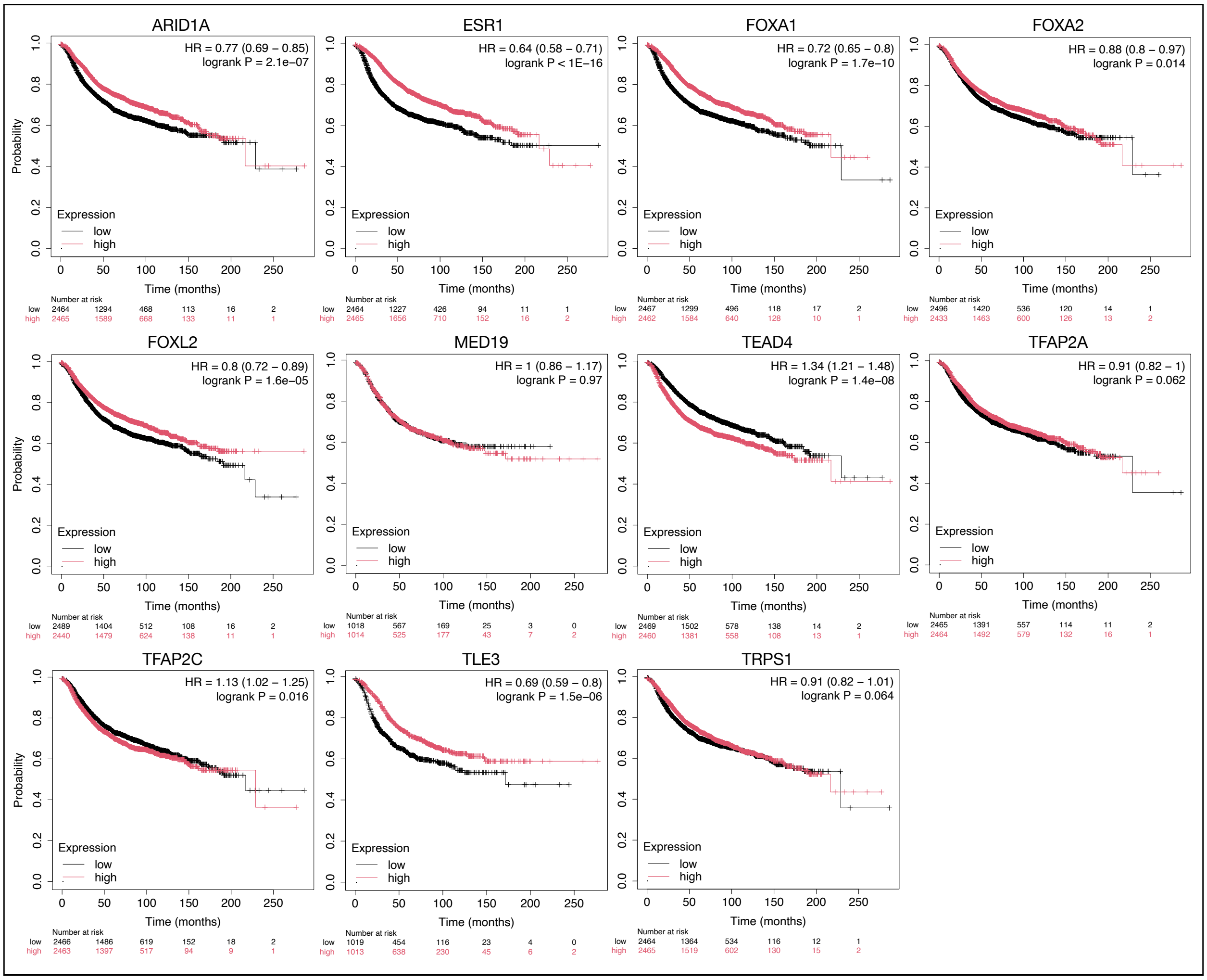


**Supplementary Figure 6. Kaplan-Meier plot of breast cancer-type-specific high-confidence TRs.** The Kaplan-Meier plots assess the association between gene expression levels of breast cancer-specific high-confidence TRs (ARID1A, ESR1, FOXA1, FOXA2, FOXL2, MED19, TEAD4, TFAP2A, TFAP2C, TLE3, TRPS1) and patients' survival outcomes.

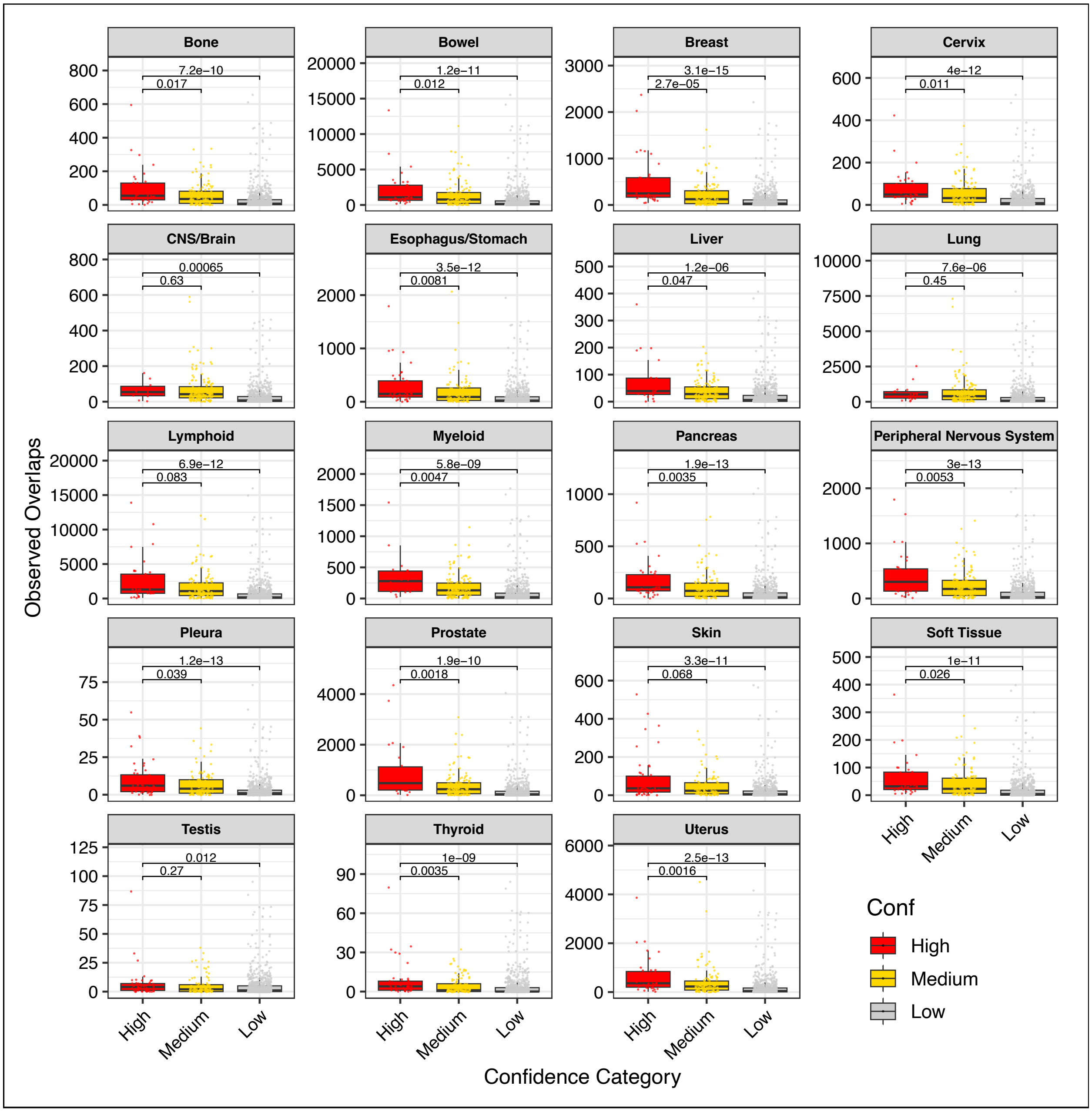


**Supplementary Figure 7. Somatic mutation sites overlapping TR binding sites.** Boxplots show the number of observed somatic mutation sites in each cancer type that overlap with TRs grouped by high-, medium-, and low-confidence categories. P-values are calculated by Wilcoxon rank sum test.

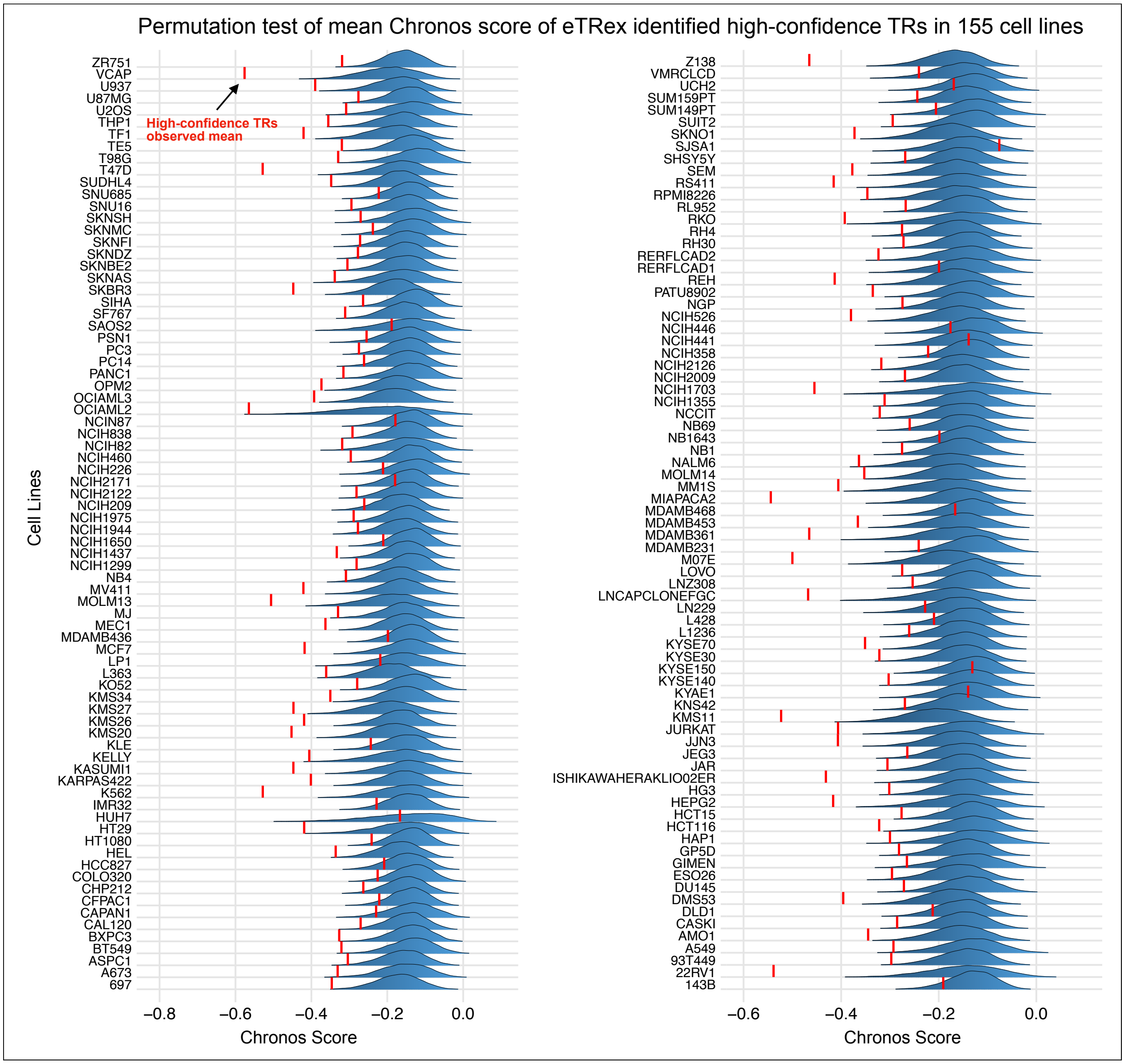


**Supplementary Figure 8. Permutation test of mean Chronos score of eTRex-identified TRs in 155 DepMap cell lines.** Conducting permutation tests of high-confidence TRs identified by eTRex in 155 cell lines that have cellular viability data in DepMap. After Benjamin-Hochberg correction, the high-confidence TRs show significant lower mean Chronos score in 121 out of 155 cell lines (adjusted p value < 0.05).

### Supplementary Note 1.

#### 1.1. Notation

| Notation | | | |
|---|---|---|---|
| $i$ | Index of TR with multiple reference datasets. | $p_{ij}(p_{i'1})$ | The probability of an informative "bin" being matching in the $j^{\text{th}}$ reference dataset for the $i^{\text{th}}$ ($i'^{\text{th}}$) TR ($j \equiv 1$ for the $i'^{\text{th}}$ TR). |
| $i'$ | Index of TR with a single reference dataset. | $\theta_{ij}(\theta_{i'1})$ | Probit transformed $p_{ij}$ ($p_{i'1}$). Dataset level importance score. |
| $j$ | Index of reference dataset. | $\theta_i(\theta_{i'})$ | The TR-level importance score. |
| $k$ | Index of informative bin. | $\sigma_i^2$ | Variance across different datasets of the $i^{th}$ TR. |
| $J_i$ | Total number of reference datasets of the $i^{th}$ TR. | $\sigma_0^2$ | Variance for TRs with only one dataset. |
| $\mathcal{M}$ | Total number of TRs with multiple datasets. | $\mu$ | Global mean of TR-level importance. |
| $\mathcal{M}^c$ | Total number of TRs with a single dataset. | $\tau^2$ | Global variability of TR-level importance. |
| $x_{ijk}(x_{i'1k})$ | Binary status of the $k^{\text{th}}$ informative bin of the $j^{\text{th}}$ reference dataset for the $i^{\text{th}}$ ($i'^{\text{th}}$) TR ($j \equiv 1$ for the $i'^{\text{th}}$ TR), with 1 being matching and 0 being mismatching | $L_\mu, U_\mu$ | Lower and upper bound of uniform prior of $\mu$. |
| $n_{ij}(n_{i'1})$ | Number of "informative" bins when matching input with the $j^{th}$ reference dataset for the $i^{\text{th}}$ ($i'^{\text{th}}$) TR ($j \equiv 1$ for the $i'^{\text{th}}$ TR). | $a, b$ | Tiny positive values of Gamma parameters for priors of $1/\tau^2$, $1/\sigma_0^2$, and $1/\sigma_i^2$. |
| $z_{ijk}$ ($z_{i'1k}$) | Auxiliary latent variable follows Gaussian distribution centered at $\theta_{ij}$ ($\theta_{i'1}$) with unit variance. | | |

#### 1.2. eTRex model

eTRex is a Bayesian hierarchical probit model that leverages the strategy of data augmentation and implements variational inference to improve computational efficiency. We index TRs with multiple reference ChIP-seq datasets by $i = 1,2,\ldots,\mathcal{M}$, each having $j = 1,2,\ldots,J_i$ datasets. We further index each informative bin when comparing the input to the $j^{\text{th}}$ reference dataset of the $i^{\text{th}}$ TR by $k = 1,2,..,n_{ij}$, where $n_{ij}$ denotes the number of informative bins. Similarly, TRs with a single reference dataset are indexed by $i' = 1,2,\ldots,\mathcal{M}^c$, all with $j \equiv 1$ dataset, and their informative bins are indexed as $k = 1,2,\ldots,n_{i'1}$.

Let $x_{ijk}$ be the binary status of the $k^{\text{th}}$ informative bin of the $j^{\text{th}}$ reference dataset for the $i^{\text{th}}$ TR, where $x_{ijk} = 1$ indicates "matching" and $x_{ijk} = 0$ indicates "mismatching". The probability of an informative bin being "matching" is denoted by $p_{ij}$ ($p_{i'1}$), with $P(x_{ijk} = 1) = p_{ij}$ ($P(x_{i'1k} = 1) = p_{i'1}$). Let $\theta_{ij} = \Phi^{-1}(p_{ij})$ $(\theta_{i'1} = \Phi^{-1}(p_{i'1}))$ be the probit transformed $p_{ij}(p_{i'1})$, which is interpreted as the importance score measuring the consistency of binding patterns between input and reference at the dataset level. We further integrate information from multiple datasets of the same TR to get the importance score at the TR level ($\theta_i/\theta_{i'}$) by assuming $\theta_{ij} \sim N(\theta_i, \sigma_i^2)$ and $\theta_{i'1} \sim N(\theta_{i'}, \sigma_0^2)$; all $\theta_i(\theta_{i'})$ are governed by a global importance score $\mu$ and variance $\tau^2$ by assuming $\theta_i, \theta_{i'} \sim N(\mu, \tau^2)$. In this way, for each TR with multiple datasets, we model its dataset-level consistency scores using a normal distribution with distinctive mean $\theta_i$ and variance $\sigma_i^2$, which reflect its overall consistency and within-TR heterogeneity, respectively. TRs with only one dataset share a common variance $\sigma_0^2$ for information pooling.

The prior distributions over the hyperparameters follow standard choices in Bayesian hierarchical modeling: we place a weakly-informative prior on the global mean, that is $\mu \sim N(0, \tau_0^2)$, and assign gamma priors based on empirical knowledge learned from data to the precision terms $1/\tau^2 \sim Gamma(a, b)$ and $1/\sigma_i^2 \sim Gamma(a, b)$ for $i = 0,1, \ldots, \mathcal{M}$. Under these specifications, the full hierarchical model can be written as:

$$x_{ijk} \sim Bernoulli(p_{ij}), x_{i'1k} \sim Bernoulli(p_{i'1})$$

$$\theta_{ij} = \Phi^{-1}(p_{ij}), \theta_{i'1} = \Phi^{-1}(p_{i'1})$$

$$\theta_{ij} \sim N(\theta_i, \sigma_i^2), \theta_{i'1} \sim N(\theta_{i'}, \sigma_0^2)$$

$$\theta_i, \theta_{i'} \sim N(\mu, \tau^2)$$

Notably, there is no conjugate prior exist for the probit transformed variable $\theta_{ij}$. To overcome this problem, we leverage the data augmentation strategy by introducing a group of latent variables $z_{ijk}$ and $z_{i'1k}$. These latent variables follow Gaussian distribution centered at $\theta_{ij}$ ($\theta_{i'1}$), and $x_{ijk}$ ($x_{i'1k}$) become deterministic conditional on the sign of $z_{ijk}$ ($z_{i'1k}$).

$$z_{ijk} \sim N\left(\theta_{ij}, 1\right), \qquad z_{i'1k} \sim N(\theta_{i'1}, 1)$$

$$x_{ijk} | z_{ijk} = \begin{cases} 1, if\ z_{ijk} > 0 \\ 0, if\ z_{ijk} < 0 \end{cases}$$

$$x_{i'1k} | z_{i'1k} = \begin{cases} 1, if\ z_{i'1k} > 0 \\ 0, if\ z_{i'1k} < 0 \end{cases}$$

Let the observed data be $X = \left\{ \{x_{ijk}\}_{i=1,j=1,k=1}^{M,J_i,n_{ij}}, \{x_{i'1k}\}_{i'=1,k=1}^{M_c,n_{i'1}} \right\}$. The unknown quantities, including parameters and latent variables involved in the eTRex model, are collectively denoted by $\Theta = \left\{ \mu, \tau^2, \sigma_0^2, \{\theta_i\}_{i=1}^{M}, \{\theta_{i'}\}_{i'=1}^{M_c}, \{\sigma_i^2\}_{i=1}^{M}, \{\theta_{ij}\}_{i=1,j=1}^{M,J_i}, \{\theta_{i'1}\}_{i'=1}^{M_c}, \{z_{ijk}\}_{i=1,j=1,k=1}^{M,J_i,n_{ij}}, \{z_{i'1k}\}_{i'=1,k=1}^{M_c,n_{i'1}} \right\}$. The full probability model of eTRex is then given by the product of all prior and conditional distributions:

$$p(X, \Theta) = p(\mu)p(\theta)p(\tau^2) \prod_{i=1}^{\mathcal{M}} p(\theta_i | \mu, \tau^2) p\left(\sigma_i^2\right) \prod_{i'=1}^{\mathcal{M}^c} p(\theta_{i'} | \mu, \tau^2) \prod_{i=1}^{\mathcal{M}} \prod_{j=1}^{J_i} p\left(\theta_{ij} \middle| \theta_i, \sigma_i^2\right)$$

$$\times \prod_{i'=1}^{\mathcal{M}^c} p(\theta_{i'1} | \theta_{i'}, \sigma_0^2) \prod_{i=1}^{\mathcal{M}} \prod_{j=1}^{J_i} \prod_{k=1}^{n_{ij}} p(x_{ijk} | z_{ijk}) p(z_{ijk} | \theta_{ij}) \prod_{i'=1}^{\mathcal{M}^c} \prod_{k=1}^{n_{i'1}} p(x_{i'1k} | z_{i'1k}) p(z_{i'1k} | \theta_{i'1})$$

#### 1.3. Coordinate Ascent Mean-field Variational Inference (MFVI)

eTRex implements a variational inference algorithm that recasts the problem of estimating the joint posterior distribution as an optimization problem: it seeks to minimize the KL divergence between the chosen variational distribution family and the true posterior distribution. eTRex uses a mean-field variational family which assumes that the latent variables and parameters are mutually independent, and each governed by a distinct factor in the variational distribution $q(\Theta)$,

$$q(\Theta) = q(\mu)q(\tau^2)q(\sigma_0^2)\prod_{i=1}^{\mathcal{M}} q(\sigma_i^2)\prod_{i=1}^{\mathcal{M}} q(\theta_i)\prod_{i'=1}^{\mathcal{M}^c} q(\theta_{i'})\prod_{i=1}^{\mathcal{M}}\prod_{j=1}^{J_i} q(\theta_{ij})\prod_{i'=1}^{\mathcal{M}^c} q(\theta_{i'1})$$

$$\times \prod_{i=1}^{\mathcal{M}}\prod_{j=1}^{J_i}\prod_{k=1}^{K_{ij}} q(z_{ijk})\prod_{i'=1}^{\mathcal{M}^c}\prod_{k=1}^{K_{i'1}} q(z_{i'1k}).$$

Within the mean-field variational family, CAVI iteratively updates each factor while holding all other factors fixed. The optimal solution of a factor has the general form,

$$\ln\big(q(\theta)\big) = E_{\Theta/\theta}\big[\ln\big(p(\mathrm{X}, \Theta)\big)\big] + \mathrm{const},$$

where the expectation is taken with respect to the current variational distributions of all remaining latent variables or parameters. For instance, based on the joint distribution $p(X, \Theta)$ and factorized variational distribution $q(\Theta)$. To update the factor $q(z_{ijk})$ and $q(z_{i'1k})$, the optimal solution can be written as

$$\ln\Big(q\big(z_{ijk}\big)\Big) = E_{\Theta\backslash z_{ijk}}\big[\ln\big(p(\Theta, X)\big)\big] + const$$

$$= \ln\Big(p\big(x_{ijk}\big|z_{ijk}\big)\Big) + E_{q(\theta_{ij})}\Big[\ln\Big(p\big(z_{ijk}\big|\theta_{ij}\big)\Big)\Big] + const.$$

(1) Update factors $q(z_{ijk})$ and $q(z_{i'1k})$:

Given $p\big(x_{ijk}\big|z_{ijk}\big) = 1\big(z_{ijk} \geq 0\big)^{x_{ijk}} 1\big(z_{ijk} < 0\big)^{1-x_{ijk}}$,

$$q^*\big(z_{ijk}\big) = \begin{cases} TN^+\big(z_{ijk}\big|\mu_{ij}, 1\big), & if\ x_{ijk} = 1 \\ TN^-\big(z_{ijk}\big|\mu_{ij}, 1\big), & if\ x_{ijk} = 0 \end{cases}.$$

Similarly, we have

$$q^*(z_{i'1k}) = \begin{cases} TN^+(z_{i'1k}|\mu_{i'1}, 1), & if\ x_{i'1k} = 1 \\ TN^-(z_{i'1k}|\mu_{i'1}, 1), & if\ x_{i'1k} = 0' \end{cases}$$

where $\mu_{ij} = E_{q(\theta_{ij})}[\theta_{ij}]$ and $\mu_{i'1} = E_{q(i'1)}[\theta_{i'1}]$.

(2) Update factor $q(\theta_{ij})$ and $q(\theta_{i'1})$:

$$\ln\left(q^*(\theta_{ij})\right) = E_{q(\Theta\backslash\theta_{ij})}\left[\ln\left(p\left(\theta_{ij}\middle|\theta_i,\sigma_i^2\right)\right)\right] + \sum_{k=1}^{n_{ij}} E_{q(z_{ijk})}\left[\ln\left(p\left(z_{ijk}\middle|\theta_{ij}\right)\right)\right] + const$$

$$= -\frac{1}{2}E_{q(\sigma_i^2)}\left[\frac{1}{\sigma_i^2}\right]\theta_{ij}^2 + E_{q(\theta_i,\sigma_i^2)}\left[\frac{\theta_i}{\sigma_i^2}\right]\theta_{ij} - \frac{n_{ij}\theta_{ij}^2}{2} + \sum_{k=1}^{n_{ij}} E_{q(z_{ijk})}\left[z_{ijk}\right]\theta_{ij} + const$$

$$= -\frac{E_{q(\sigma_i^2)}\left[\frac{1}{\sigma_i^2}\right] + n_{ij}}{2}\theta_{ij}^2 + \left(E_{q(\theta_i)}[\theta_i]E_{q(\sigma_i^2)}\left[\frac{1}{\sigma_i^2}\right] + \sum_{k=1}^{n_{ij}} E_{q(z_{ijk})}\left[z_{ijk}\right]\right)\theta_{ij} + const.$$

Therefore,

$$q^*(\theta_{ij}) \sim N(m_{ij}, S_{ij}),$$

$$S_{ij} = \left(E_{q(\sigma_i^2)}\left[\frac{1}{\sigma_i^2}\right] + n_{ij}\right)^{-1}, m_{ij} = S_{ij}\left(E_{q(\theta_i)}[\theta_i]E_{q(\sigma_i^2)}\left[\frac{1}{\sigma_i^2}\right] + \sum_{k=1}^{n_{ij}} E_{q(z_{ijk})}\left[z_{ijk}\right]\right).$$

Similarly,

$$q^*(\theta_{i'1}) \sim N(m_{i'1}, S_{i'1}),$$

$$S_{i'1} = \left(E_{q(\sigma_0^2)}\left[\frac{1}{\sigma_0^2}\right] + n_{i'1}\right)^{-1}, m_{i'1} = S_{i'1}\left(E_{q(\theta_{i'})}[\theta_{i'}]E_{q(\sigma_0^2)}\left[\frac{1}{\sigma_0^2}\right] + \sum_{k=1}^{n_{i'1}} E_{q(z_{i'1k})}[z_{i'1k}]\right).$$

(3) Update factor $q(\theta_i)$ and $q(\theta_{i'})$:

$$q^*(\theta_i) = -\frac{E_{q(\tau^2)}\left[\frac{1}{\tau^2}\right] + J_i E_{q(\sigma_i^2)}\left[\frac{1}{\sigma_i^2}\right]}{2}\theta_i^2 + \left(E_{q(\tau^2)}\left[\frac{1}{\tau^2}\right]E_{q(\mu)}[\mu] + \sum_{j=1}^{J_i} E_{q(\theta_{ij})}\left[\theta_{ij}\right]E_{q(\sigma_i^2)}\left[\frac{1}{\sigma_i^2}\right]\right)\theta_i.$$

Thus,

$$q^*(\theta_i) \sim N(T_i, V_i),$$

$$V_i = \left(E_{q(\tau^2)}\left[\frac{1}{\tau^2}\right] + J_i E_{q(\sigma_i^2)}\left[\frac{1}{\sigma_i^2}\right]\right)^{-1},$$

$$T_i = V_i\left(E_{q(\tau^2)}\left[\frac{1}{\tau^2}\right]E_{q(\mu)}[\mu] + \sum_{j=1}^{J_i} E_{q(\theta_{ij})}[\theta_{ij}]E_{q(\sigma_i^2)}\left[\frac{1}{\sigma_i^2}\right]\right).$$

Similarly,

$$q^*(\theta_{i'}) \sim N(T', V'),$$

$$V_{i'} = \left(E_{q(\tau^2)}\left[\frac{1}{\tau^2}\right] + E_{q(\sigma_0^2)}[\frac{1}{\sigma_0^2}]\right)^{-1},$$

$$T_{i'} = V_{i'}\left(E_{q(\tau^2)}\left[\frac{1}{\tau^2}\right]E_{q(\mu)}[\mu] + E_{q(\theta_{i'1})}[\theta_{i'1}]E_{q(\sigma_0^2)}\left[\frac{1}{\sigma_0^2}\right]\right).$$

(4) Update factor $q(\mu)$ and variance terms:

To make the update derivation easier, as gamma update is more commonly used in MFVI, we denote $\omega = \frac{1}{\tau^2}$, and similarly for $\omega_i = \frac{1}{\sigma_i^2}$ and $\omega_0 = \frac{1}{\sigma_0^2}$, which are corresponding to the precisions. It is also noted that the terms $E\left[\frac{1}{\tau^2}\right] = E[\omega]$, $E\left[\frac{1}{\sigma_0^2}\right] = E[\omega_0]$, and $E\left[\frac{1}{\sigma_i^2}\right] = E[\omega_i]$ in this reparameterization. Thus, given that the variance terms follow inverse-gamma prior, we have $\omega, \omega_0, \omega_1 \dots, \omega_{\mathcal{M}}$ follow gamma distribution $\omega, \omega_0, \omega_1, \dots \omega_{\mathcal{M}} \sim Gamma(a, b)$. Also, we have $\mu \sim N(0, 10^2)$. Therefore, we have

$$q^*(\mu) \sim N(\mu^*, D),$$

$$D = \left(\frac{1}{\tau_0^2} + I E_{q(\tau^2)}\left[\frac{1}{\tau^2}\right]\right)^{-1},$$

$$\mu^* = \left(I + \frac{1}{\tau_0^2 E_{q(\tau^2)}\left[\frac{1}{\tau^2}\right]}\right)^{-1} \left(\sum\nolimits_{i=1}^{M} E_{q(\theta_i)}[\theta_i] + \sum\nolimits_{i'=1}^{M^c} E_{q\left(\theta_{i'}\right)}[\theta_{i'}]\right),$$

$$q^*(\omega) \sim Gamma(a^*, b^*),$$

$$a^* = a + \frac{I}{2},$$

$$b^* = b + \frac{1}{2}\left(IE_{q(\mu)}[\mu^2] - 2E_{q(\mu)}[\mu]\left(\sum_{i=1}^{M} E_{q(\theta_i)}[\theta_i] + \sum_{i'=1}^{M^c} E_{q\left(\theta_{i'}\right)}\left[\theta_{i'}\right]\right) + \sum_{i=1}^{M} E_{q(\theta_i)}[\theta_i^2] + \sum_{i'=1}^{M^c} E_{q\left(\theta_{i'}\right)}\left[\theta_{i'}^2\right]\right),$$

$$q^*(\omega_0) \sim Gamma(a_0^*, b_0^*),$$

$$a_0^* = a + \frac{M^c}{2},$$

$$b_0^* = b + \frac{1}{2}\sum_{i'=1}^{M^c} \left(E_{q(\theta_{i'1})}[\theta_{i'1}^2] - 2E_{q(\theta_{i'1})}[\theta_{i'1}]E_{q(\theta_{i'})}[\theta_{i'}] + E_{q(\theta_{i'})}[\theta_{i'}^2]\right),$$

$$q^*(\omega_i) \sim Gamma(a_i^*, b_i^*),$$

$$a_i^* = a + \frac{J_i}{2},$$

$$b_i^* = b + \frac{1}{2}\sum_{j=1}^{J_i} \left(E_{q(\theta_{ij})}[\theta_{ij}^2] - 2E_{q(\theta_{ij})}[\theta_{ij}]E_{q(\theta_i)}[\theta_i] + E_{q(\theta_i)}[\theta_i^2]\right).$$

Also, expectation terms can be derived and given by:

$$E\left[z_{ijk}\right] = \begin{cases} \mu_{ij} + \phi_{ij}/(1-\Phi_{ij}), & if\ x_{ijk} = 1 \\ \mu_{ij} - \phi_{ij}/(\Phi_{ij}), & if\ x_{ijk} = 0 \end{cases},$$

$$E[z_{i'1k}] = \begin{cases} \mu_{i'1} + \phi_{i'1}/(1-\Phi_{i'1}), & if\ x_{i'1k} = 1 \\ \mu_{i'1} - \phi_{i'1}/(\Phi_{i'1}), & if\ x_{i'1k} = 0 \end{cases},$$

$$E\left[\theta_{ij}\right] = m_{ij}, \qquad E[\theta_{i'1}] = m_{i'1},$$

$$E[\theta_i] = T_i, \qquad E[\theta_{i'}] = T_i',$$

$$E[\mu] = \mu^*, \qquad E\left[\frac{1}{\tau^2}\right] = E[\omega] = \frac{a^*}{b^*},$$

$$E\left[\frac{1}{\sigma_0^2}\right] = E[\omega_0] = \frac{a_0^*}{b_0^*}, \qquad E\left[\frac{1}{\sigma_i^2}\right] = E[\omega_i] = \frac{a_i^*}{b_i^*},$$

$$E[\mu^2] = \mu^{*2} + D,$$

$$E\left[\theta_i^2\right] = T_i^2 + V_i,$$

$$E\left[\theta_{i'}^2\right] = T_{i'}^2 + V_{i'},$$

$$E\left[\theta_{ij}^2\right] = m_{ij}^2 + S_{ij},$$

$$E\left[\theta_{i'1}^2\right] = m_{i'1}^2 + S_{i'1},$$

where $\phi_{ij} = \phi(-\mu_{ij})\ (\phi_{i'1} = \phi(-\mu_{i'1}))$ and $\Phi_{ij} = \Phi(-\mu_{ij})\ (\Phi_{i'1} = \Phi(-\mu_{i'1}))$. Based on the updating equations (1)-(4), we have:

In the $t^{th}$ iteration:

(1) $z_{ijk}$ & $z_{i'1k}$

$$\mu_{ij}^{(t)} = m_{ij}^{(t-1)}, \mu_{i'1}^{(t)} = m_{i'1}^{(t-1)},$$

$$E^{(t)}\left[z_{ijk}\right] = \begin{cases} \mu_{ij}^{(t)} + \phi_{ij}^{(t)}/(1-\Phi_{ij}^{(t)}), & if\ x_{ijk} = 1 \\ \mu_{ij}^{(t)} - \phi_{ij}^{(t)}/(\Phi_{ij}^{(t)}), & if\ x_{ijk} = 0 \end{cases},$$

$$E^{(t)}[z_{i'1k}] = \begin{cases} \mu_{i'1}^{(t)} + \phi_{i'1}^{(t)}/(1-\Phi_{i'1}^{(t)}), & if\ x_{i'1k} = 1 \\ \mu_{i'1}^{(t)} - \phi_{i'1}^{(t)}/(\Phi_{i'1}^{(t)}), & if\ x_{i'1k} = 0 \end{cases}.$$

(2) $\theta_{ij}$ & $\theta_{i'1}$

$$E^{(t)}\left[\theta_{ij}\right] = m_{ij}^{(t)} = \underbrace{\left(\frac{a_i^{*\,(t-1)}}{b_i^{*(t-1)}} + n_{ij}\right)^{-1}}_{S_{ij}^{(t)}} \left(T_i^{(t-1)} \frac{a_i^{*\,(t-1)}}{b_i^{*(t-1)}} + \sum_{k=1}^{n_{ij}} E^{(t)}\left[z_{ijk}\right]\right),$$

$$E^{(t)}[\theta_{i'1}] = m_{i'1}^{(t)} = \underbrace{\left(\frac{a_0^{*\,(t-1)}}{b_0^{*(t-1)}} + n_{i'1}\right)^{-1}}_{S_{i'1}^{(t)}} \left(T_{i'}^{(t-1)} \frac{a_0^{*\,(t-1)}}{b_0^{*(t-1)}} + \sum_{k=1}^{n_{i'1}} E^{(t)}[z_{i'1k}]\right),$$

$$E^{(t)}\left[\theta_{ij}^2\right] = {m_{ij}^{(t)}}^2 + S_{ij}^{(t)},$$

$$E^{(t)}\left[\theta_{i'1}^2\right] = {m_{i'1}^{(t)}}^2 + S_{i'1}^{(t)}.$$

(3) $\theta_i$ & $\theta_{i'}$

$$E^{(t)}[\theta_i] = T_i^{(t)} = \underbrace{\left(\frac{a^{*(t-1)}}{b^{*(t-1)}} + J_i \frac{a_i^{*\,(t-1)}}{b_i^{*(t-1)}}\right)^{-1}}_{V_i^{(t)}} \left(\frac{a^{*(t-1)}}{b^{*(t-1)}} \mu^{*(t-1)} + \sum_{j=1}^{J_i} m_{ij}^{(t)} \frac{a_i^{*\,(t-1)}}{b_i^{*(t-1)}}\right),$$

$$E^{(t)}[\theta_{i'}] = T_{i'}^{(t)} = \underbrace{\left(\frac{a^{*(t-1)}}{b^{*(t-1)}} + \frac{a_0^{*\,(t-1)}}{b_0^{*(t-1)}}\right)^{-1}}_{V_{i'}^{(t)}} \left(\frac{a^{*(t-1)}}{b^{*(t-1)}} \mu^{*(t-1)} + m_{i'1}^{(t)} \frac{a_0^{*\,(t-1)}}{b_0^{*(t-1)}}\right),$$

$$E^{(t)}\left[\theta_i^2\right] = (T_i^{(t)})^2 + V_i^{(t)},$$

$$E^{(t)}\left[\theta_{i'}^2\right] = (T_{i'}^{(t)})^2 + V_{i'}^{(t)}.$$

(4) $\mu$

$$E^{(t)}[\mu] = \mu^{*(t)} = \left( I + \frac{1}{\tau_0^2 \frac{a^*}{b^*}} \right)^{-1} \left( \sum\nolimits_{i=1}^{M} T_i^{(t)} + \sum\nolimits_{i'=1}^{M^c} T_{i'}^{(t)} \right),$$

$$E^{(t)}[\mu^2] = \mu^{*(t)} + D^{(t)} = \left( I + \frac{1}{\tau_0^2 \frac{a^*}{b^*}} \right)^{-1} \left( \sum\nolimits_{i=1}^{M} T_i^{(t)} + \sum\nolimits_{i'=1}^{M^c} T_{i'}^{(t)} \right) + \left( \frac{1}{\tau_0^2} + I \frac{a^{*(t-1)}}{b^{*(t-1)}} \right)^{-1}.$$

(5) $\tau^2$

$$E^{(t)}\left[\frac{1}{\tau^2}\right] = \frac{a^{*(t)}}{b^{*(t)}},$$

$$a^{*(t)} = a + \frac{I}{2},$$

$$b^{*(t)} = b + \frac{1}{2}\left( I\left(\mu^{*(t)} + D^{(t)}\right) - 2\mu^{*(t)} \left( \sum_{i=1}^{M} T_i^{(t)} + \sum_{i'=1}^{M^c} T_{i'}^{(t)} \right) + \sum_{i=1}^{M} \left( (T_i^{(t)})^2 + V_i^{(t)} \right) \right.$$
$$\left. + \sum_{i'=1}^{M^c} \left( (T_{i'}^{(t)})^2 + V_{i'}^{(t)} \right) \right).$$

(6) $\sigma_0^2$

$$E^{(t)}\left[\frac{1}{\sigma_0^2}\right] = \frac{a_0^{*(t)}}{b_0^{*(t)}},$$

$$a_0^{*(t)} = a_0 + \frac{M^c}{2},$$

$$b_0^{*(t)} = b_0 + \frac{1}{2} \sum_{i'=1}^{M^c} \left( {T_{i'}^{(t)}}^2 + V_{i'}^{(t)} - 2 m_{i'1}^{(t)} T_{i'}^{(t)} + {m_{i'1}^{(t)}}^2 + S_{i'1}^{(t)} \right).$$

(7) $\sigma_i^2$

$$E^{(t)}\left[\frac{1}{\sigma_i^2}\right] = \frac{a_i^{*(t)}}{b_i^{*(t)}},$$

$$a_i^{*(t)} = a_i + \frac{M}{2},$$

$$b_i^{*(t)} = b_i + \frac{1}{2}\sum_{j=1}^{J_i}\left(T_i^{(t)^2} + V_i^{(t)} - 2T_i^{(t)}m_{ij}^{(t)} + m_{ij}^{(t)^2} + S_{ij}^{(t)}\right).$$

By now, we have all the updating equations in one iteration.
